# H2A.Z deposition by SWR1C involves multiple ATP-dependent steps

**DOI:** 10.1101/2022.01.11.475888

**Authors:** Jiayi Fan, Andrew T. Moreno, Alexander S. Baier, Joseph J. Loparo, Craig L. Peterson

## Abstract

The histone variant H2A.Z is a conserved feature of nucleosomes flanking protein-coding genes. Deposition of H2A.Z requires ATP-dependent replacement of nucleosomal H2A by a chromatin remodeler related to the multi-subunit enzyme, yeast SWR1C. How these enzymes use ATP to promote this nucleosome editing reaction remains unclear. Here we use single-molecule and ensemble methodologies to identify three ATP-dependent phases in the H2A.Z deposition reaction. Real-time analysis of single nucleosome remodeling events reveals an initial, priming step that occurs after ATP addition that likely involves transient DNA unwrapping from the nucleosome. Priming is followed by rapid loss of histone H2A, which is subsequently released from the H2A.Z nucleosomal product. Surprisingly, the rates of both priming and the release of the H2A/H2B dimer are sensitive to ATP concentration. This complex reaction pathway provides multiple opportunities to regulate the timely and accurate deposition of H2A.Z at key genomic locations.

## Introduction

The eukaryotic genome is packaged into the nucleoprotein structure called chromatin, which at its most fundamental level is composed of nucleosomes. The nucleosome consists of two copies of histones H2A, H2B, H3, and H4 wrapped by ∼147bp of DNA (Luger et al., 1997). All essential nuclear processes including transcription, DNA replication and repair require dynamic regulation of chromatin structure by post-translational histone modifications, ATP-dependent chromatin remodeling, and the exchange of canonical histones for their variants (Clapier et al., 2017).

One key histone variant is histone H2A.Z, which replaces canonical H2A in a replication-independent manner at the nucleosomes proximal to gene transcription start sites and transcriptional enhancers, DNA double-stranded breaks, and replication origins (Albert et al., 2007; Barski et al., 2007; Raisner et al., 2005), as H2A.Z is believed to be a major regulator of gene expression (Subramanian* et al., 2015), DNA repair (Adkins et al., 2013; Ye et al., 2012), and replication origin licensing (Long et al., 2019), respectively. H2A.Z is evolutionarily conserved from yeast to human, and it is indispensable for fly (Daal and Elgin, 1992) and mammalian embryonic development (Faast et al., 2001). H2A.Z is deposited by the megadalton complexes of the INO80 subfamily chromatin remodelers, characterized by SWR1C in yeast (Mizuguchi et al., 2004) and its orthologs Tip60/p400(Gévry et al., 2007) and SRCAP (Ruhl et al., 2006) in mammals. Mutations that disrupt these H2A.Z-depositing machines lead to abnormal dosage and localization of H2A.Z on chromatin and can result in a variety of diseases, such as Floating-Harbor Syndrome (Greenberg et al., 2019; Hood et al., 2012; Reschen et al., 2012), uterine leiomyoma (Berta et al., 2021), and cancer (Hsu et al., 2018; Mattera et al., 2009; Slupianek et al., 2010).

Chromatin remodeling enzymes have been separated into four distinct subfamilies – SWI/SNF, ISWI, CHD, and INO80 (Clapier et al., 2017). Most chromatin remodelers, including SWR1C, initiate their remodeling activity by binding to superhelical location 2 (SHL 2) of nucleosomal DNA, located 20 base pairs from the center of nucleosomal symmetry, known as the dyad. Prior DNA-histone crosslinking and single-molecule Förster resonance energy transfer (smFRET) experiments have shown that remodelers of the SWI/SNF, ISWI, and CHD subfamilies bind to one DNA strand of SHL2 and induce a ∼1-2-nucleotide shift(Li et al., 2019; Winger et al., 2018; Yan and Chen, 2020). This shift along the presumed tracking strand is then reiterated on the guide strand, leading to a ∼1-2bp DNA translocation toward the dyad upon ATP binding and hydrolysis, synchronizing the ATP cycle to the stepwise translocation of nucleosomal DNA during remodeler sliding. Currently, it is unknown whether the INO80 subfamily performs a similar DNA translocation cycle.

SWR1C stands out among remodelers for utilizing ATP hydrolysis to not slide DNA, but to exchange a nucleosomal H2A/H2B dimer with a H2A.Z/H2B dimer (Anand et al., 2015). Previous *in vitro* studies using bulk assays have shown that SWR1C sequentially deposits two H2A.Z-H2B dimers on a substrate that mimics the +1 nucleosome adjacent to the nucleosome-free region (NFR) with an asymmetric preference for NFR-distal exchange (Luk et al., 2010; Singh et al., 2019). However, bulk assays cannot resolve the precise timing of the two exchanges occurring on the same nucleosome or detect reaction intermediates that occur prior, during, and after exchange, and whether they are sensitive to ATP. Furthermore, we do not know how the different phases of the ATP hydrolysis cycle impact the interactions of SWR1C with the nucleosome during dimer exchange.

Here, we address these mechanistic questions of SWR1C remodeling by employing single-base pair and single-nucleosome resolution techniques of site-specific DNA-histone crosslinking and single-molecule Förster Resonance Energy Transfer (smFRET) microscopy, respectively. Using crosslinking, we show that SWR1C does not change the path of DNA on the histone octamer surface during its ATP-dependent deposition of H2A.Z. Instead, single-molecule experiments show that dimer exchange occurs through three distinct phases, two of which are sensitive to ATP concentrations. Finally, we use a fluorescence polarization assay to show that ATP hydrolysis dramatically weakens nucleosome binding, consistent with ATP-dependent release of SWR1C following H2A.Z deposition, as suggested by our smFRET studies. Together, these comprehensive assays construct an intricate picture of SWR1C dimer exchange and provide extensive insights on how H2A.Z deposition is regulated on a molecular level.

## Results

### SWR1C does not alter the path of nucleosomal DNA during binding or H2A.Z deposition

Previous ensemble studies of H2A.Z deposition have suggested that SWR1C uses ATP hydrolysis to transiently unwrap DNA from the nucleosomal edge, followed by stepwise replacement of the two H2A/H2B dimers (Singh et al., 2019). How SWR1C promotes DNA unwrapping is not clear. Many remodeling enzymes use a cycle of ATP binding and hydrolysis to translocate the ATPase subunit along one strand of DNA in a 3’ to 5’ direction, leading to movement of DNA on the histone octamer surface (Clapier et al., 2017). One simple possibility is that SWR1C may use such movement to weaken histone-DNA interactions, leading to DNA unwrapping. For other remodelers, this translocation reaction has been monitored using nucleosomal substrates with histones harboring site-specific, photo-activatable crosslinking agents, such as 4-azidophenacyl bromide (APB) (Hota and Bartholomew, 2012; Winger et al., 2018). For many remodelers, such as Isw1 and Chd1, binding of the enzyme to nucleosomes is sufficient to induce a 1-2bp movement of one strand of DNA (Li et al., 2019; Winger et al., 2018). This remodeler-induced perturbation of the DNA path is often sensitive to whether the remodeler is bound to a non-hydrolyzable ATP analog, which is thought to represent an intermediate in the translocation reaction (Bowman and Deindl, 2019; Yan and Chen, 2020).

To investigate whether SWR1C can alter the path of nucleosomal DNA, a center-positioned nucleosome was reconstituted that harbored an engineered APB-modified cysteine residue on histone H2B-Q59C. One end of the ‘601’ nucleosome positioning sequence was labeled with a Cy5 fluorophore, and the other end was labeled with a FAM fluorophore to visualize DNA-histone crosslinks from each nucleosome face (Figure 1A). Consistent with previous work (Winger et al., 2018), APB modification of H2B-Q59C leads to a predominant UV-induced DNA crosslink ±53nt from the nucleosomal dyad (SHL±5.5) (Figure 1B and Supplementary Figure S1). Addition of either the Chd1 or ISW2 remodeler led to a new crosslink at +55bp, and ISW2 also induced a new crosslink at the opposite nucleosome face at position -55bp (Figure 1B and Supplementary Figure S1A,C). This is consistent with 2bp movement of DNA from the entry/exit point of the nucleosome towards the nucleosomal dyad, as previously shown (Winger et al., 2018). A more dramatic alteration in DNA crosslinks was observed after addition of the RSC remodeler and AMP-PNP (Supplementary Figure S1B). In contrast, addition of SWR1C had no detectable impact on DNA crosslinks at the +53 or -53 position, and the crosslink pattern in the presence of SWR1C was not altered by further addition of ADP or non-hydrolyzable ATP analogs, AMP-PNP or ADP•BeF_3_^-^ (Figure 1B and Supplementary Figure S1C). Nucleosomes were also reconstituted with the APB group on H2B-S93C that monitors DNA at SHL3.0, only one helical turn from where the Swr1 ATPase binds to nucleosomal DNA (Figure 1A). Addition of ISW2 to the H2B-S93C-APB nucleosome produced a new crosslink at +33/+34, consistent with DNA translocation, but addition of SWR1C had no impact on the crosslinking pattern (Supplementary Figure S1F,G).

**Figure 1.**
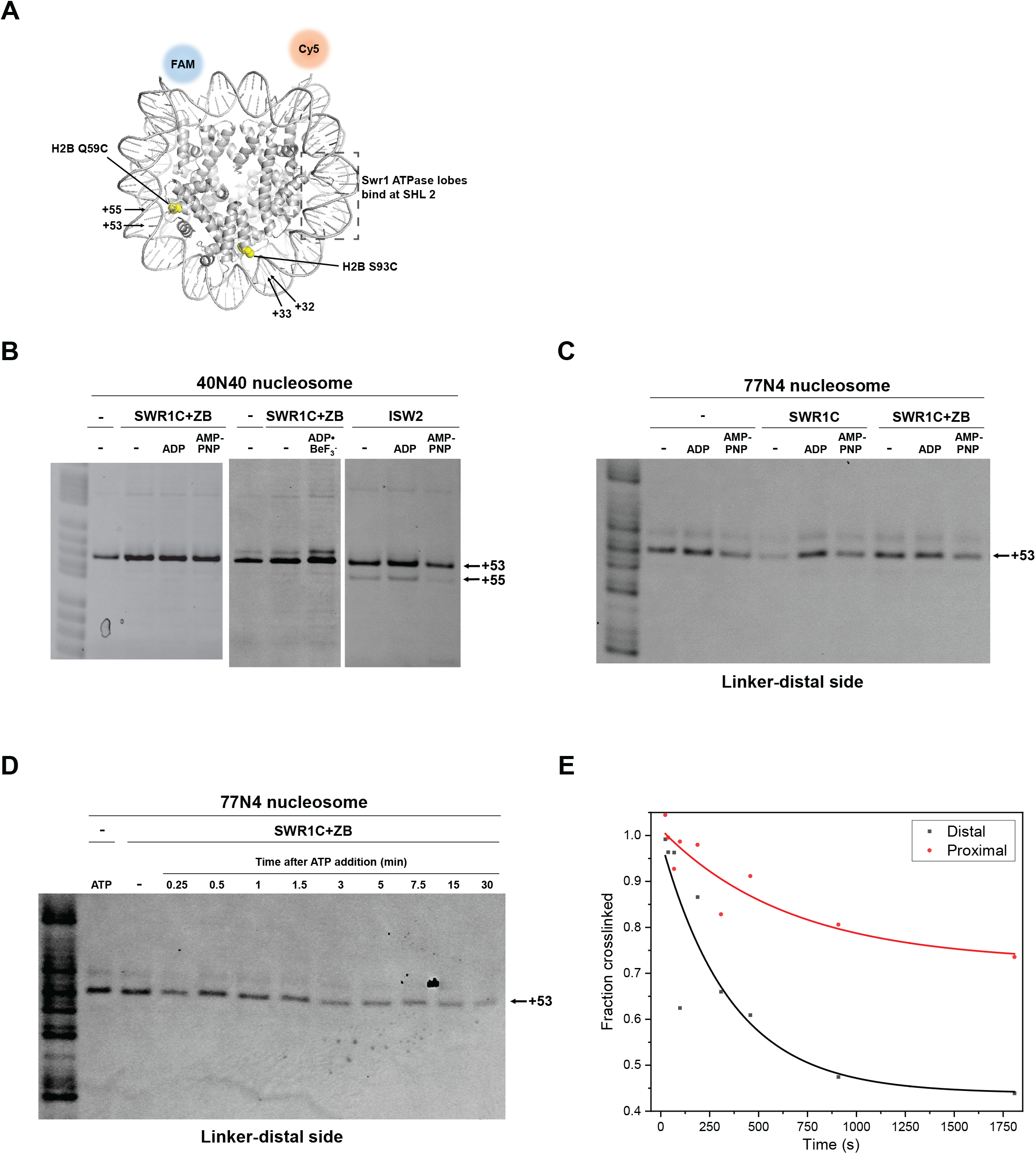
Site-directed DNA-histone mapping shows that SWR1C does not change the path of nucleosomal DNA in any nucleotide state as a part of its ATP-dependent dimer exchange activity. (A) DNA-histone mapping schematics on a yeast nucleosome (PDB: 1ID3). APB-labeled H2B-Q59C generates a DNA crosslink at SHL5.5 (+53) while H2B-S93C generates a crosslink at SHL3 (+32). (B) ISW2 binding in apo or ADP-bound state induces a 2-nucleotide translocation at SHL5.5 toward the dyad on a center-positioned nucleosome, while SWR1C binding does not, regardless of the presence of nucleotides or H2A.Z/H2B dimer. (C) SWR1C binding does not alter the nucleosomal DNA path at SHL5.5 of an asymmetric nucleosome template. (D) SWR1C dimer exchange is robust in the absence of DNA translocation under 3 μM ATP. (E) Quantification of SWR1C dimer exchange time course for the DNA linker-distal (Figure 1D) or linker-proximal (Figure S1H) nucleosomal H2A/H2B dimer, capturing SWR1C preference for distal dimer exchange.

In our previous study, end-positioned nucleosomal substrates were designed with 77bp of flanking linker DNA so that it might reflect the asymmetry of a promoter-proximal nucleosome located next to a nucleosome free region (NFR) (Singh et al., 2019). On this substrate, SWR1C preferentially exchanges the H2A/H2B dimer that is distal to the long linker, consistent with the pattern of H2A.Z deposition *in vivo* (Rhee et al., 2014). Both orientations of an end-positioned nucleosome (77N4 and 4N77) were reconstituted with histones harboring H2B-Q59C, and APB crosslinking was performed in the absence or presence of SWR1C (Figure 1C and Supplementary Figure S1D,E). For both substrates, SWR1C did not induce changes in the crosslinking pattern adjacent to either the distal or proximal H2A/H2B dimer interfaces (Figure 1C and Supplementary Figure S1D,E). Furthermore, addition of H2A.Z/H2B dimer and nucleotides to the SWR1C binding reactions had no impact.

To ensure that SWR1C was active on these APB-modified substrates, H2A.Z/H2B dimers and a low concentration of ATP (3 μM) were added to binding reactions, and a crosslinking time-course was performed (Figure 1D,E; see also Supplementary Figure S1H). Addition of ATP led to a time-dependent loss of H2B-DNA crosslinks, consistent with dimer exchange. Furthermore, loss of crosslinks for the linker-distal dimer interface occurred more rapidly than the linker-proximal interface, consistent with preferential and asymmetric replacement of the linker-distal dimer. Notably, no changes in the crosslinking positions were detected during the dimer exchange reaction. Together, these data suggest that SWR1C may be distinct from other remodelers, and that this remodeler may not induce substantial changes in the path of nucleosomal DNA on the histone octamer surface during H2A.Z deposition. Furthermore, the data are consistent with previous studies demonstrating that SWR1C induces transient unwrapping from the nucleosomal edge (Singh et al., 2019; Willhoft et al., 2018).

### Single-molecule FRET detects preferential and sequential dimer exchange

To probe the dynamics of the H2A.Z deposition reaction in real-time on individual nucleosomes, we developed a smFRET SWR1C dimer exchange assay. An end-positioned nucleosome was reconstituted on a nucleosome positioning sequence harboring a biotin moiety on the end of a 117bp free DNA linker with an ATTO 647N acceptor fluorophore located on the 4bp linker of the opposite end. Histone H2A was labeled with a Cy3B donor fluorophore at an engineered cysteine residue, placing the Cy3B and ATTO 647N fluorophores at the appropriate distance to function as a FRET pair (Figure 2A). Following immobilization of biotinylated nucleosomes on a streptavidin-coated slide, nucleosomes were imaged by total internal reflection fluorescence (TIRF) microscopy. The nucleosomal substrate is similar to that used for ensemble dimer exchange reactions, in which SWR1C activity leads to an ATP-dependent loss of FRET as the Cy3B-labeled dimer is replaced with unlabeled H2A.Z (Singh et al., 2019). Since the labeling efficiency of H2A is only ∼ 80%, nucleosome reconstitutions show three clusters of FRET values – nucleosomes with Cy3B on only the linker-distal dimer have a high FRET efficiency (∼0.78-0.9), nucleosomes with Cy3B on only the linker-proximal dimer have a low FRET efficiency (∼0.4-0.57), and nucleosomes with both labeled dimers have an intermediate FRET value (∼0.58-0.77) (see Figure 2D, right panels; see also Figure 3A-E for individual, nucleosome trajectories). Consequently, the smFRET assay can report on the timing and efficiency of dimer eviction events from each nucleosome face on individual, surface-immobilized nucleosomes.

**Figure 2.**
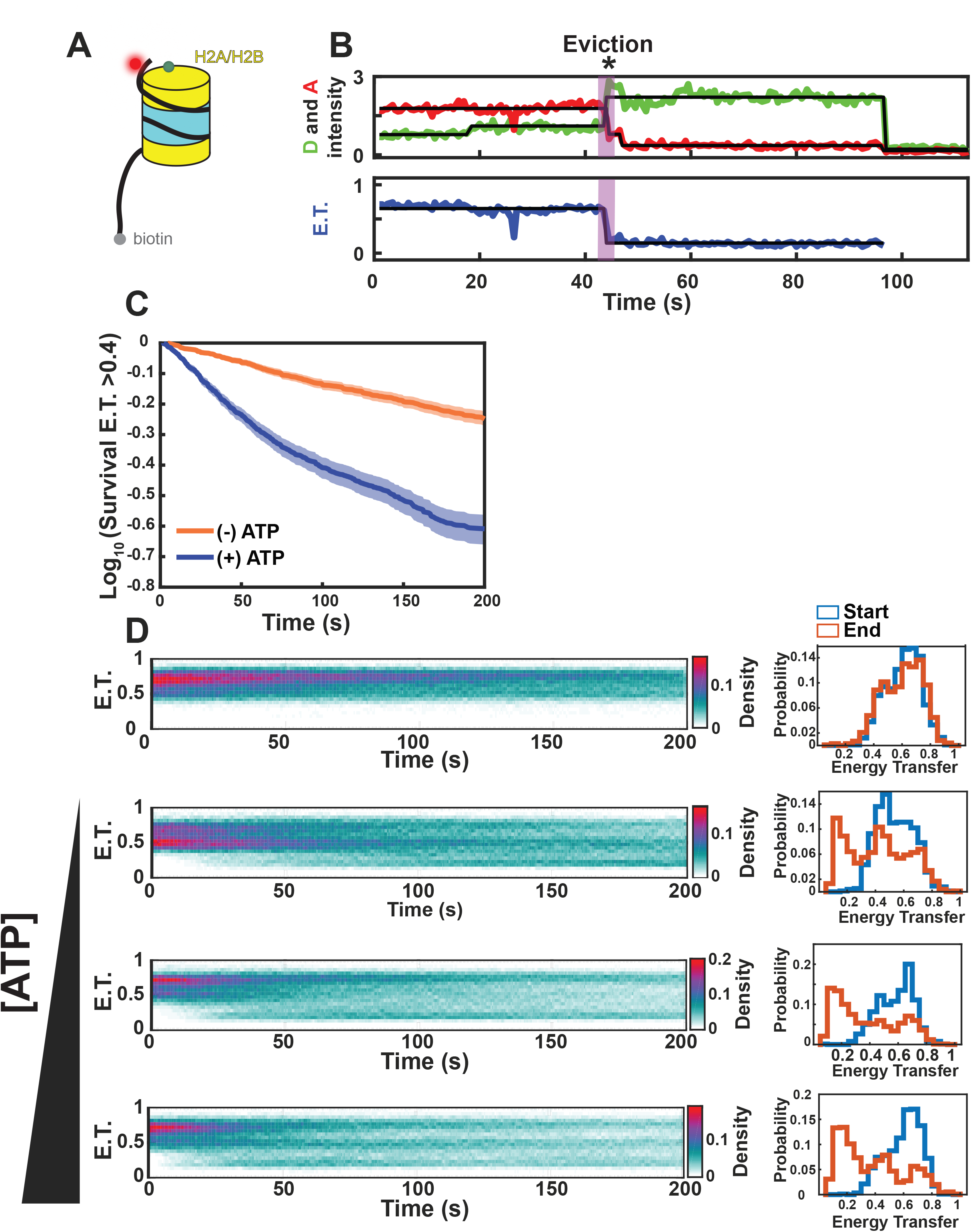
SWR1C eviction of H2A from the nucleosome is ATP dependent. (A) Schematic of nucleosome substrate for smFRET studies. Mononucleosomes contained 4bp and 117bp DNA linkers, with an ATTO 647N fluorophore positioned on the end of the short linker and a biotin group on the long linker. Histone H2A was labeled at the C-terminal domain with a Cy3B fluorophore. Only a single, labelled H2A is shown for simplicity. (B) Example trajectory. The Cy3B donor fluorophore was excited, and donor emission (green) and ATTO 647N acceptor emission (red) were recorded (top panel) and used to calculate energy transfer efficiency (blue, bottom panel). The eviction of H2A is marked by purple triangle. (C) Energy transfer (>0.4) survival kinetics (Kaplan-Meier estimate) for 0 μM ATP (N=962) (orange) and 100 μM ATP (N=939) (blue). The x axis indicates dwell time in the high E.T. state (solid line). Shaded areas, 95% confidence intervals. (D) right-side: Time-resolved energy transfer histograms for tethered nucleosomes with increasing ATP concentrations (0, 0.5, 5, 100 μM) left-side: histogram of the FRET distribution at the start (blue) and end (red) of the trajectory.

**Figure 3.**
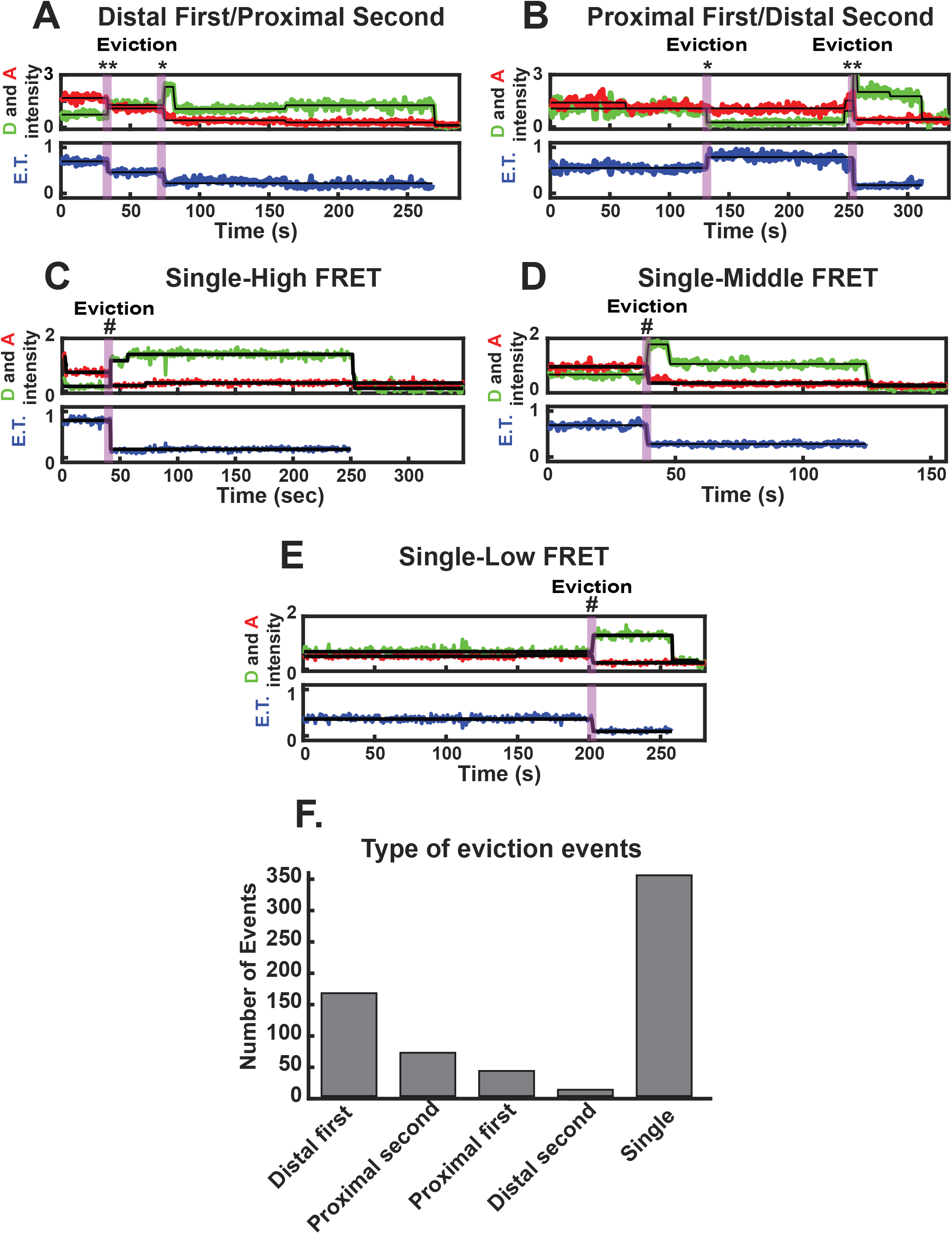
SWR1C evicts distal and proximal H2A from the nucleosome. (A-E) Example trajectory. The Cy3B donor fluorophore was excited, and donor emission (green) and ATTO 647N acceptor emission (red) were recorded (top panel) and used to calculate energy transfer efficiency (blue, bottom panel). (A) Example trajectory of SWR1C evicting distal H2A (**) from the nucleosome followed by the eviction of proximal H2A (*). (B) Example trajectory of SWR1C evicting proximal H2A (*) from the nucleosome followed by the eviction of distal H2A (**). (C) Example trajectory of SWR1C eviction of H2A (#) from a nucleosome in a high energy transfer state. (D) Example trajectory of SWR1C eviction of H2A (#) from a nucleosome in a medium energy transfer state. (E) Example trajectory of SWR1C eviction of H2A (#) from the nucleosome in a low energy transfer state. (F) Observed events for each type of eviction (A-E) in the presence of 100 μM ATP (N=939). Single eviction represents summation of events C-E.

The dimer exchange reaction was reconstituted by preincubating SWR1C with nucleosomes and a ∼2-fold molar excess of H2A.Z/H2B dimers and the H2A.Z-specific histone chaperone, Chz1 before tethering the substrates in the flow cell (Luk et al., 2007). Addition of ATP initiated the exchange reaction, which resulted in the frequent loss of FRET on individual nucleosomal substrates consistent with eviction of the donor-labeled H2A/H2B dimer (Figure 2B,C). The dynamics of the various FRET states are readily observed by plotting the FRET efficiency as a function of time (Figure 2D). These kymographs show loss of the initial FRET population (Energy Transfer [E.T.] > 0.5) and the appearance of a low FRET population (E.T. ∼0.2). In addition, the rate of these transitions is dependent on ATP concentration (Figure 2D). Eviction events were defined as persistent loss of FRET from an initial state greater than E.T. = 0.4 to a final FRET state less than E.T. = 0.35. While loss of FRET was sometimes observed in the absence of ATP, these events are likely due to photobleaching, and the inclusion of nucleotide greatly increased the number of apparent dimer eviction events (ATP 0 μM = 0.036 min^-1^, ATP 100 μM = 0.48 min^-1^).

Given that there are two H2A/H2B dimers per nucleosome, and the labeling of H2A was sub-stoichiometric, five types of eviction events are anticipated (Figure 3A-E). For nucleosomes harboring two labeled H2A/H2B dimers (Figure 3A,B), there was a preferential, ATP-dependent decrease to a low FRET value, consistent with asymmetric exchange of the linker-distal dimer, as previously observed by an ensemble FRET assay (Figure 3A,F; (Singh et al., 2019)). Importantly, 40% of these nucleosomes also showed a second decrease in FRET, consistent with a stepwise exchange of the two H2A/H2B dimers (Figure 3A,B,F and Figure S3). We also observed a population of nucleosomes where the two dimers appeared to be evicted in a single step (Figure 3E,F).

### smFRET detects multiple ATP-dependent steps during H2A.Z deposition

Analysis of individual FRET trajectories showed that the H2A.Z deposition reaction had three distinct phases (Figure 4A). Following addition of ATP, nucleosomes exhibited a “priming” period prior to a stable loss of FRET. Priming is ATP-mediated, as the duration of the priming phase (t_prime_) decreased with increasing ATP concentration (Figure 4B). For instance, in the presence of 0.5 μM ATP the lifetime of the priming phase was 92 (82.8-102.9 95% C.I.) seconds and decreased to 55 (51.0-59.6 95% C.I.) seconds with 100 μM ATP (Figure 4B and Supplementary Table S1). Following the priming phase, the loss of FRET occurred rapidly. The length of time for eviction (t_evict_) was ∼2-3s, and it was largely insensitive to ATP concentration (Figure 4C,E). After rapid eviction, the H2A-H2B dimer remained associated with the H2A.Z nucleosome, with complete loss from the immobilized nucleosome occurring over a longer period (t_release_). Surprisingly, the half-life of t_release_ decreased with increasing ATP concentration, 219 (204.5-240 95% C.I.) seconds at 0.5 μM ATP and 96 (68.0-120.5 95% C.I.) seconds at 100 μM ATP, indicating that the release of H2A/H2B from the final product is an ATP-dependent step (Figure 4D, Supplementary Figure S3, and Supplementary Table S1).

**Figure 4.**
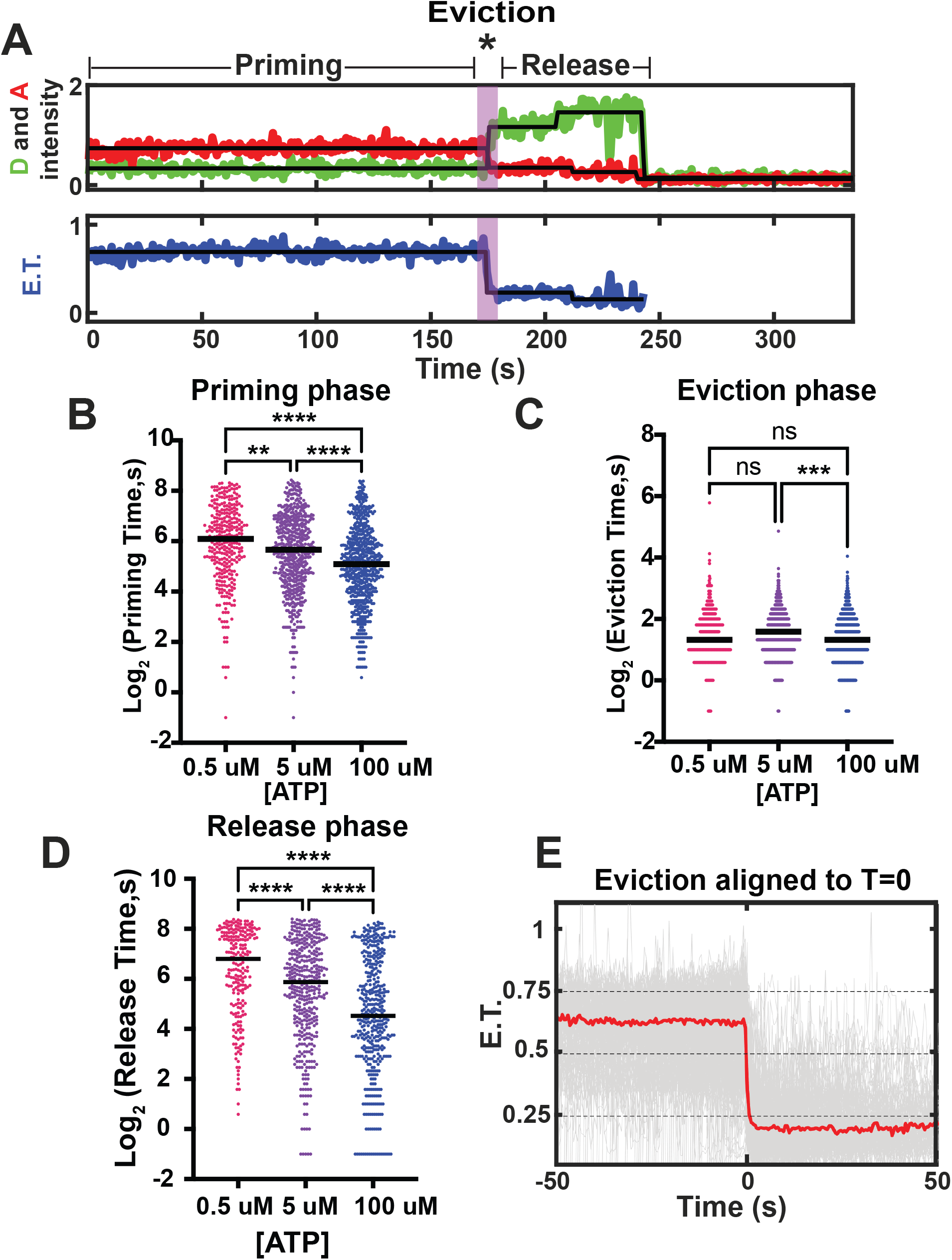
The duration of the priming and release phase is dependent on ATP concentration. (A) Example trajectory highlighting priming, eviction, and release intervals. The Cy3B donor fluorophore was excited, and donor emission (green) and ATTO 647N acceptor emission (red) were recorded (top panel) and used to calculate energy transfer efficiency (blue, bottom panel). (B) Distribution of the priming phase duration with the indicated ATP concentration. (C) Distribution of the eviction phase duration with the indicated ATP concentration. (D) Distribution of the release phase duration with the indicated ATP concentration. In (B-D) black line is the median, ns, not significant, **, p < 0.0014, ***, p < 0.004, ****, p < 0.0001 for one-way ANOVA Tukey’s multiple comparison test. (E) Alignment of E.T. trajectories with eviction phase centered at zero with 100 μM ATP (N=635), red line is the median.

Further analysis of the initial, priming phase of the H2A.Z deposition reaction revealed a large number of reversible transitions between high and low FRET states (Figure 5A,B). Fluctuations in FRET were scored as reversible if the transitions initiated from an E.T. state > 0.6 to an E.T. state < 0.45 and returning to the initial state. These transitions increased in frequency and duration after ATP addition (Figure 5B,C). In the absence of ATP, we observed reversible FRET transitions in 55/491 trajectories (1.2 min^-1^), with an average dwell time of 1.49 ± 2.32 seconds, while in the presence of ATP, we observed 100/487 trajectories (2.1 min^-1^) having FRET changes with a dwell time of 3.4 ± 5.8 seconds (Supplementary Table S2). Given their reversible nature, these FRET transitions likely correspond to transient nucleosome unwrapping events (Singh et al., 2019; Willhoft et al., 2018).

**Figures 5.**
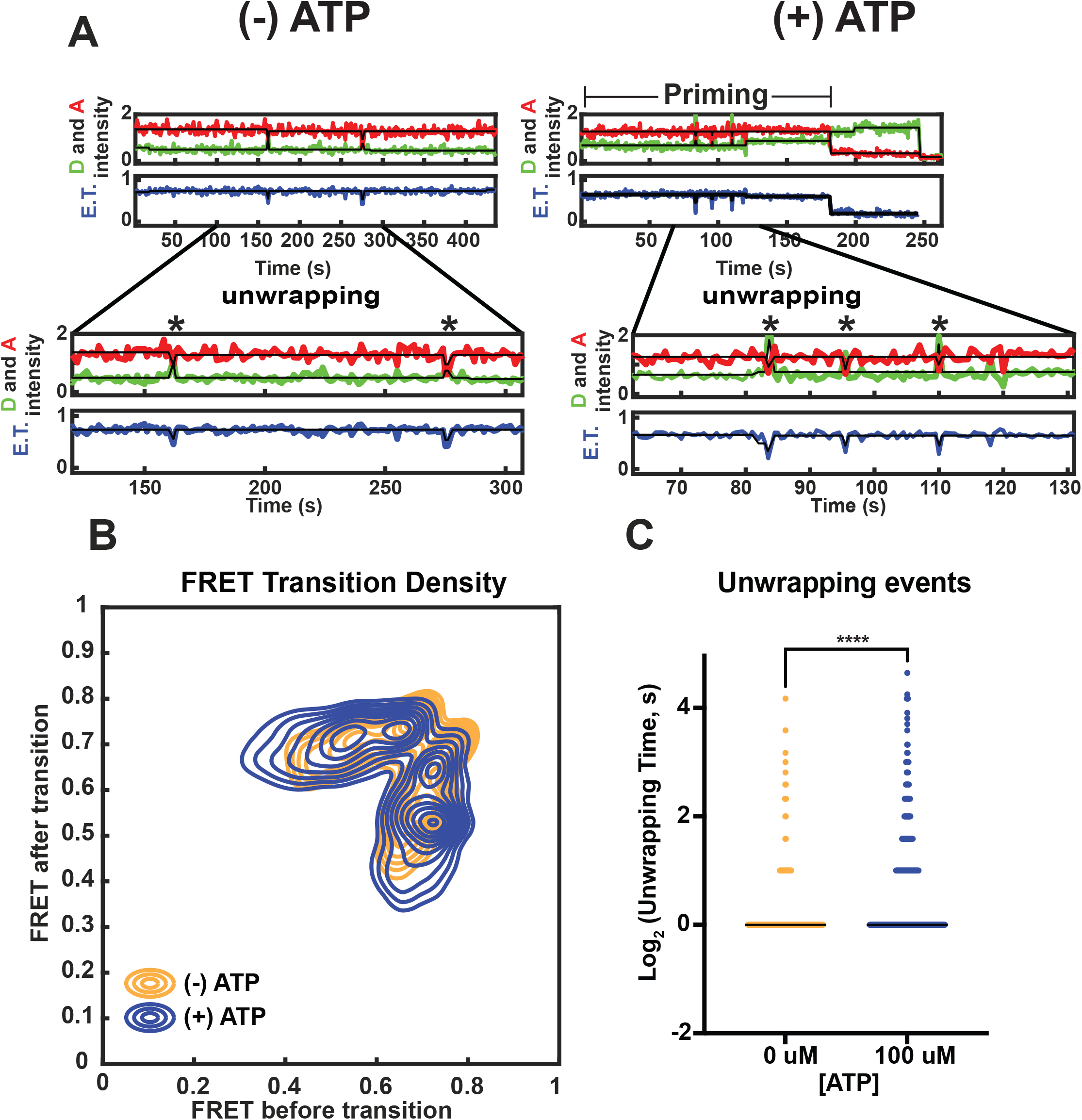
The frequency of SWR1C induced nucleosome unwrapping increases with the addition of ATP. (A) Example trajectory highlighting unwrapping events (*) that occur in the priming phase for 0 μM ATP (left side) and 100 μM ATP (N=487). The Cy3B donor fluorophore was excited, and donor emission (green) and ATTO 647N acceptor emission (red) were recorded (top panel) and used to calculate energy transfer efficiency (blue, bottom panel). (B) Priming phase E.T. transition density plot. The intensity of E.T. transitions are normalized to the total observation window for 0 μM ATP (N=491) (orange) and 100 μM ATP (N=487) (blue). (C) Distribution in unwrapping event dwell times for 0 μM ATP (N=491) (orange) and 100 μM ATP (N=487) (blue), ****, p < 0.0001 for unpaired student t-test.

To directly monitor deposition of H2A.Z/H2B dimers in real-time, nucleosomes were reconstituted with ATTO 647N-labeled DNA and unlabeled histone octamers, while free H2A.Z/H2B dimers were labeled on H2A.Z with Cy3B (Supplementary Figure S4). Nucleosomes were first incubated with SWR1C and immobilized on a streptavidin-coated slide. The deposition reaction was then initiated with the addition of Cy3B-H2A.Z/H2B dimers and ATP. Trajectories of Cy3B-H2A.Z fluorescence at individual nucleosomes showed association of single or at times two or more Cy3B-H2A.Z/H2B dimers (Supplementary Figure S4). This simultaneous association of multiple dimers might reflect the ability of SWR1C to interact with more than one dimer on the nucleosome. Control regions of interest that lacked nucleosomes showed far fewer Cy3B-H2A.Z binding events (Supplementary Figure S4B) confirming that most binding is nucleosome specific. Nearly all nucleosomes (N=307) exhibited binding (89%) of at least one Cy3B-H2A.Z dimer over the course of the 5-minute movie (Supplementary Figure S4). Cy3B-H2A.Z/H2B dimers rapidly colocalized with immobilized nucleosomes. For instance, we observed Cy3B-H2A.Z/H2B dimers binding to 50% of the nucleosomes within 10 seconds (Supplementary Figure S4C). The interaction between a Cy3B-H2A.Z/H2B dimer and SWR1C-nucleosome was relatively stable with a mean lifetime of 34 seconds (N = 1,015), although some individual Cy3B-H2A.Z/H2B dimers remained associated for hundreds of seconds (Supplementary Figure S4, S5, and Supplementary Table 3). However, as many of the tethered nucleosomes showed multiple dimers of Cy3B-H2A.Z/H2B simultaneously associated (Supplementary Figure S5), the average lifetime of individual dimers may be shorter. After colocalization, a subset of trajectories exhibited an increase in FRET, likely due to successful deposition of H2A.Z (Supplementary Figure S4A and S5). Importantly, the length of time at which co-localized Cy3B-H2A.Z/H2B was incorporated into the nucleosome was similar to the duration of the priming phase for H2A eviction (Supplementary Figure S4A, S5, and Supplementary Table 3), consistent with the anticipated concerted eviction and deposition reaction (Luk et al., 2010; Singh et al., 2019).

### Binding of SWR1C to nucleosomes is sensitive to nucleotides

Real-time analyses of the H2A.Z deposition reaction indicated that a replaced H2A/H2B dimer is released from an immobilized nucleosome in an ATP-dependent reaction. Previous work has implicated the Swc5 subunit of SWR1C as a candidate histone chaperone that binds to the released H2A/H2B dimer and facilitates eviction and replacement (Sun and Luk, 2017). One possibility is that the ATP-dependent loss of H2A/H2B from nucleosomes reflects the release of a SWR1C-H2A/H2B complex. This model suggests that the binding of SWR1C to either an H2A- or H2A.Z-containing nucleosome may be regulated by ATP binding or hydrolysis. To test this hypothesis, a ATTO 647N-labeled, 77N4 nucleosome was used in fluorescence polarization assays to quantify the nucleosome binding affinity of SWR1C. In the absence of nucleotides, SWR1C bound an H2A nucleosome with an apparent K_d_ of 12 nM ± 2 nM. Strikingly, addition of ATP led to a large decrease in binding affinity to an H2A-containing nucleosome (31 nM ± 8 nM), and a decrease in binding was also observed with ADP (21 nM ± 4 nM, but not AMP-PNP (14 nM ± 4 nM) (Figure 6B). Together, these data indicate the strength of SWR1C-nucleosome interactions is modulated during the ATP cycle and hydrolysis may function to release the enzyme following H2A.Z deposition.

**Figure 6.**
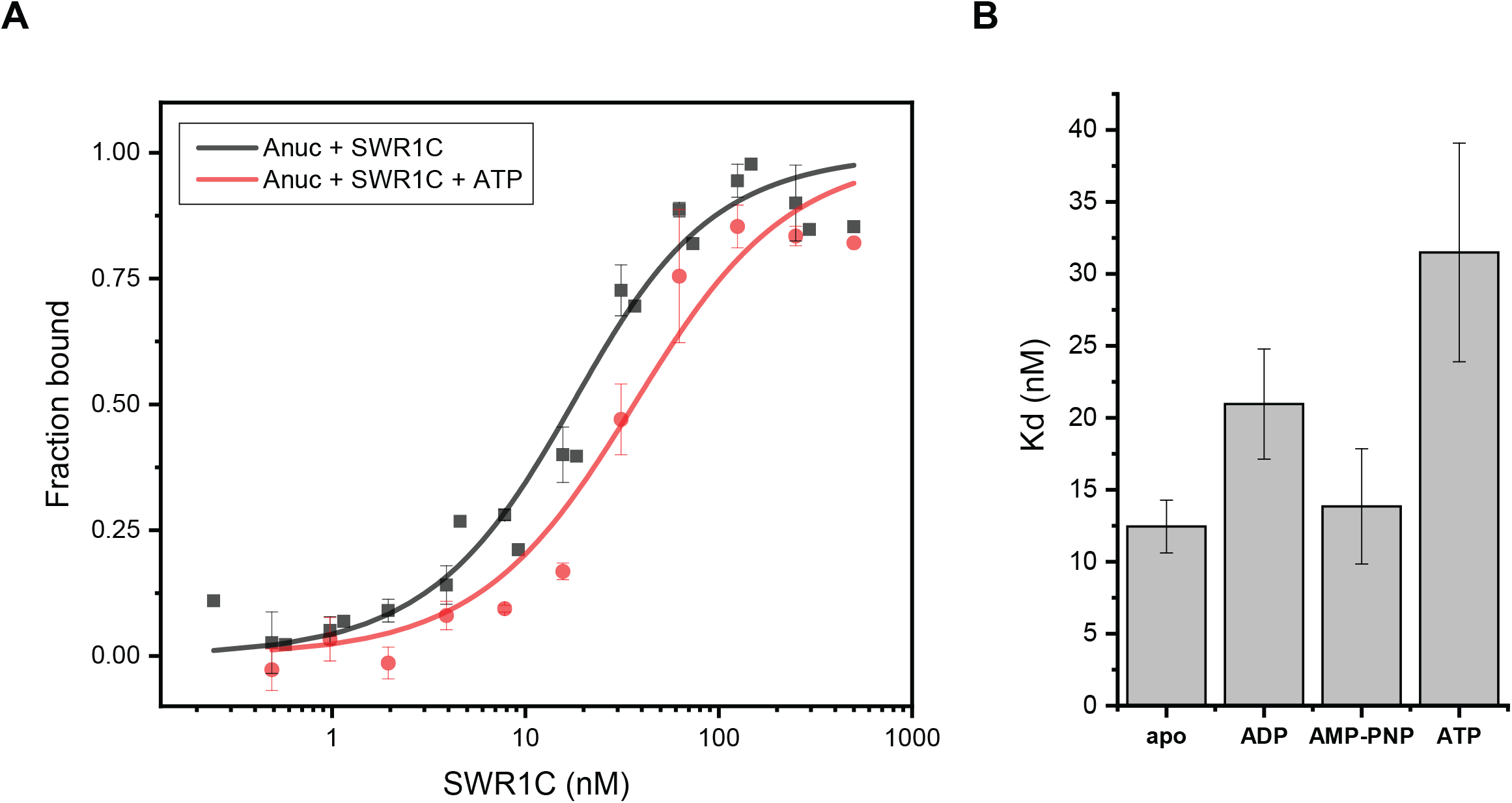
Fluorescence polarization assays show that ATP hydrolysis promotes the release of SWR1C from the H2A-nucleosome. (A) Normalized fluorescence polarization plots and fitted curves for SWR1C binding to H2A-nucleosome without nucleotide (black) or in the presence of 1 mM ATP (red), and to H2A.Z-nucleosome in the apo state (green). Each curve represents a global fit of 3-4 replicates with the standard error indicated by the error bars. The concentration of SWR1C on the X-axis is displayed on a log_10_-scale. (B) The dissociation constant (K_d_) of SWR1C on a H2A- or H2A.Z-nucleosome in different nucleotide states with 3-4 replicates for each condition and the standard error represented by the error bars.

## Discussion

SWR1C is unique among the remodeling enzymes characterized to date, as it does not use ATP hydrolysis to alter nucleosome positioning, but rather is dedicated to the replacement of nucleosomal H2A with its variant, H2A.Z (Clapier et al., 2017). Furthermore, unlike other remodelers, the H2A.Z deposition reaction is kinetically slow even under single turnover conditions, suggesting that there may be a large number of kinetic intermediates in the reaction pathway (Singh et al., 2019). Here we use a smFRET approach to identify three ATP-dependent steps of the H2A.Z deposition reaction – an initial ‘priming’ step, followed by eviction and replacement of the nucleosomal H2A/H2B dimer, and finally the release of the H2A/H2B dimer from the nucleosomal product (see Figure 7). The dimer eviction step is quite rapid (∼4s), and this timescale is similar to a single cycle of DNA translocation by the ISWI remodeler (Blosser et al., 2009). In contrast, the initial ATP-dependent priming step is quite slow and is likely to represent the rate-limiting step for the dimer exchange reaction.

**Figure 7.**
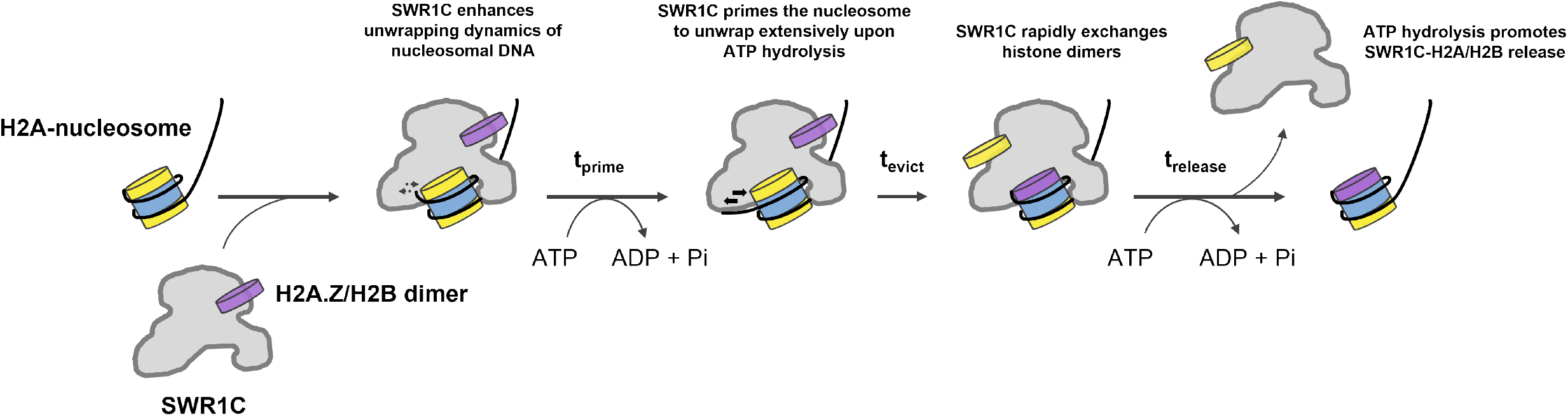
Kinetic model for H2A.Z deposition by SWR1C. Initially, SWR1C binds to the linker-distal face of an asymmetric H2A-nucleosome, such as the +1 nucleosome adjacent to the NFR. SWR1C binding at SHL2 induces stress and transient dynamics at the nucleosomal DNA edge. While nucleosome-bound, SWR1C hydrolyzes ATP for an extended period of time to prime the nucleosomal substrate for H2A eviction (t_prime_), leading to more extensive unwrapping of nucleosomal DNA. Sufficient priming allows SWR1C to rapidly perform its H2A to H2A.Z dimer exchange reaction on the nucleosome (t_evict_). The evicted H2A/H2B dimer remains associated with the SWR1C still bound to the exchanged nucleosome, until the SWR1C-H2A/H2B complex is released from the nucleosome through ATP hydrolysis (t_release_).

### The priming phase during H2A.Z deposition

Following the binding of SWR1C and H2A.Z/H2B dimers to immobilized nucleosomes and the addition of ATP, we observed a long lag phase (t_prime_) prior to eviction of the Cy3B-labeled H2A/H2B dimer. The duration of this initial priming phase decreased at higher ATP concentrations, indicating that this phase contains one or more ATP-dependent steps. A previous analysis of the ACF remodeler also identified ATP-dependent steps prior to DNA translocation (Blosser et al., 2009). However, in the case of ACF, a major component of this lag period was the ATP-sensitive binding of ACF to nucleosomes. Binding of SWR1C is unlikely to be influenced by ATP in our studies, as we first pre-incubated SWR1C with nucleosomes prior to their immobilization and ATP addition. Furthermore, our nucleosome binding assays demonstrate that ATP does not enhance the binding of SWR1C, but rather significantly weakens it. One hallmark of the initial priming phase of the H2A.Z deposition reaction is a series of FRET fluctuations that increase in frequency following ATP addition. These changes correlate well with our previous fluorescence correlation spectroscopy FRET (FCS-FRET) studies where we found that the binding of SWR1C to nucleosomes increased the dynamics of DNA-histone interactions at the nucleosomal edge by several orders of magnitude (Singh et al., 2019). These nucleosomal dynamics were then further enhanced by addition of the ATP analog, ATP**γ**S. In addition, our previous ensemble FRET studies found that SWR1C could induce transient unwrapping of DNA from the nucleosomal edge (Singh et al., 2019). We envision that repeated transient unwrapping of DNA from the nucleosomal edge may destabilize or ‘prime’ the nucleosome, lowering the kinetic barrier to the subsequent eviction of the H2A/H2B dimer.

### How is the energy of ATP hydrolysis coupled to dimer exchange?

Remodeling enzymes all contain an ATPase subunit that is a member of the large Snf2 family of DNA-stimulated ATPases (Clapier et al., 2017). The Snf2 family is part of the larger SFII superfamily of DNA/RNA helicases/translocases, and prior studies on monomeric helicases of the SFII family suggest a model whereby a remodeler ATPase cycle leads to a unidirectional, inchworm-like movement of the bi-lobular ATPase along nucleosomal DNA. Since remodelers are anchored to the nucleosomal surface by histone-binding domains, ATPase translocation can lead to movement of DNA on the octamer surface, “pulling in” DNA from the proximal, DNA entry site and propagating DNA towards the distal, exit site of the nucleosome (Clapier et al., 2017). Consistent with this model, introduction of ssDNA gaps between the nucleosomal edge and SHL2 can block the ability of remodelers to mobilize nucleosomes. Furthermore, recent analyses of the Chd1 remodeler suggests that ATP-dependent closure of the ATPase lobes is sufficient to induce a 1bp translocation step (Winger et al., 2018).

The general expectation is that SWR1C also employs a DNA translocation mechanism to drive its histone exchange reaction. Consistent with this view, the cryo-EM structure of the SWR1C-nucleosome complex contains a 1bp bulge of DNA at SHL2 (Willhoft et al., 2018). Furthermore, previous studies have shown that 2nt gaps in nucleosomal DNA can block H2A.Z deposition by SWR1C (Ranjan et al., 2015). However, unlike other remodelers that mobilize nucleosomes, only 2nt ssDNA gaps within the ATPase binding site at SHL2 block dimer exchange (±17bp to ±22bp from the nucleosomal dyad) (Ranjan et al., 2015), suggesting that the Swr1 ATPase lobes may translocate only 1-2bp of DNA. Alternatively, the gaps near SHL±2 may block SWR1C-induced deformations in DNA that precede a putative translocation step (Nodelman and Bowman, 2021). Our previous ensemble FRET studies were unable to detect translocation of DNA at the nucleosomal edge (Singh et al., 2019), and our histone-DNA crosslinking analyses presented here were also unable to detect changes in the path of nucleosomal DNA between the DNA entry point and SHL2. One possibility is that the crosslinking assay is unable to capture changes in DNA trajectories due to a technical limitation. We think this unlikely, given that this assay does detect changes in DNA-histone interactions due to the ISW2, Chd1, and RSC remodelers. Alternatively, we favor a model in which SWR1C distorts DNA at SHL2, as well as the histone octamer, without altering the register of DNA-histone contacts between the entry site and SHL2. Such a distortion would represent a high energy, strained intermediate that promotes subsequent dimer eviction and exchange. In this model, the cryo-EM structure may capture the resolution of this unstable intermediate to a stable product with fully translocated DNA.

Unlike remodelers that slide nucleosomes, such as ISWI and Chd1, SWR1C has a subunit module (Arp6/Swc6/Swc3) that binds to DNA at the nucleosomal edge and may prevent the ‘pulling’ of DNA into the nucleosome (Willhoft et al., 2018). We envision that limited translocation by the Swr1 ATPase at SHL2 acts in opposition to this barrier, distorting DNA structure without “pulling” DNA in from the entry site. This may destabilize histone-DNA contacts, leading to the transient unwrapping events that are observed during the priming stage of smFRET trajectories. A productive unwrapping event might involve stabilization of the unwrapped state by other SWR1C subunits, providing an opportunity for rapid dimer eviction and H2A.Z deposition.

### SWR1C-nucleosome binding is regulated by ATP hydrolysis

Perhaps one of the most surprising results from the smFRET analyses was the observation that release of the labeled H2A/H2B dimer from the immobilized nucleosomal product was sensitive to ATP concentration. At saturating levels of ATP, the half-life for Cy3B-H2A release was 96 seconds and when the ATP concentration was lowered to 500 nM, the lifetime increased to 219 seconds. Indeed, this last step in the H2A.Z deposition reaction was the most sensitive to ATP levels. One possibility is that the release step describes an ATP-dependent ejection of the dimer from both SWR1C and the H2A.Z product nucleosome. We favor an alternative model in which this release step reflects the ATP-dependent loss of a SWR1C-H2A/H2B complex from immobilized nucleosomes. In support of this view, we find that the affinity of SWR1C for nucleosomes is dramatically weakened by addition of ATP. Furthermore, the presence of ADP also weakened the interaction between SWR1C and the nucleosome. In contrast, the addition of the nonhydrolyzable ATP analog, AMP-PNP, had no effect on binding affinity. These data indicate a weakened interaction between SWRIC and nucleosome post ATP hydrolysis (Gangaraju et al., 2009; Ren et al., 2019). The ATP-dependent release of SWR1C from its H2A.Z nucleosomal product may facilitate re-cycling of the enzyme, promoting either a subsequent round of H2A.Z deposition on the same nucleosome or for transferring the enzyme to a new substrate. Interestingly, a recent live-cell imaging study also found that the yeast ISW1, ISW2, and Chd1 remodelers employ ATPase activity to promote fast kinetics of target search and chromatin dissociation (Kim et al., 2021).

### The stepwise asymmetry of the H2A.Z deposition reaction

From yeast to mammals, H2A.Z deposition is often targeted to the nucleosome adjacent to the start site for transcription by RNA polymerase II (Albert et al., 2007; Barski et al., 2007). Often termed the +1 nucleosome, it is inherently asymmetric, with one side flanked by a nucleosome-free region (NFR), and the other side by the +2 nucleosome (Jiang and Pugh, 2009). Our *in vitro* nucleosomal substrates mimic the asymmetry of the +1 nucleosome, as it is flanked by a 77-117 bp linker. Furthermore, DNA footprinting and cryo-EM studies have shown that interactions between SWR1C and the long linker DNA orient the ATPase lobes of the Swr1 catalytic subunit to interact with linker-distal SHL+2.0 (Anand et al., 2015; Willhoft et al., 2018), and we previously found that this leads to the preferential eviction of the linker-distal H2A/H2B dimer in an ensemble histone exchange reaction (Singh et al., 2019). High-resolution, ChIP-exo analyses of nucleosome asymmetry in yeast are consistent with asymmetric dimer exchange and enrichment of H2A.Z on the NFR-distal side of the +1 nucleosome (Rhee et al., 2014).

Two independent lines of evidence reported here are also consistent with asymmetric H2A.Z deposition. First, our histone-DNA crosslinking assays report on the ATP-dependent loss of histone H2A during the exchange reaction. The crosslinking substrates have different fluorophores at each DNA end, so this assay can determine whether a particular dimer is lost preferentially. In this case, the results mirror our previous ensemble FRET assays, supporting a biphasic loss of H2A and a preference for the linker-distal dimer. Secondly, our smFRET substrates contain H2A/H2B dimers which can each contain a fluorophore, and thus this assay can report on whether SWR1C prefers to exchange dimers located distal or proximal to the long linker DNA. Consistent with ensemble studies, we find that exchange of the linker-distal dimer is more frequent than the linker-proximal dimer. Many of the smFRET trajectories showed two, sequential H2A eviction events, and in these cases there was a clear preference for eviction of the linker-distal dimer in the first event. Interestingly, in these cases the two events were separated by a period that is similar in magnitude to only the initial priming phase, shorter than a period encompassing both priming and release intervals that is observed for single replacement events. This suggests the possibility that the two, sequential events follow a concerted pathway in which SWR1C is committed to both rounds of exchange without full release from the heterotypic H2A.Z/H2A nucleosome. Together, our data have uncovered previously unknown phases of the H2A.Z dimer exchange reaction and the unexpected requirement of ATP hydrolysis for both priming and release of the nucleosome by SWR1C, providing a new understanding of the molecular mechanism of this unique nucleosome-editing reaction.

## Acknowledgements

We thank Nathan Gioacchini (UMMS) for help with DNA sequencing analyses, Shinya Watanabe (UMMS) for the expression vector for human H3.2 C110A, and members of the Peterson and Loparo groups for helpful discussions This work was supported by grants from the National Institutes of Health [R35-GM122519] to C.L.P. and [RO1GM115487] J.J.L.

## Author Contributions

Experiments were performed by J.F. and A.T.M.; A.S.B. helped with the fluorescence polarization studies; J.F., A.T.M., C.L.P., and J.J.L. analyzed data and edited the manuscript.

## Materials and Methods

### Nucleosome Reconstitution

*S. cerevisiae* (H2A, H2A-K119C, H2A.Z, H2A.Z-K120C, H2B, H2B-Q59C, H2B-S93C), *X. laevis* (H3.1, H4), and *H. sapiens* (H3.2 C110A) histones were expressed in *E. coli* Rosetta 2(DE3)pLysS cells (except H4 which used Rosetta 2) and purified as previously described(Luger et al., 1999). The point substitutions on H2A, H2B, and H3 were generated using site-directed mutagenesis. The purified histones were reconstituted into yeast H2A/H2B-Xenopus/human H3/H4 octamers and yeast H2A.Z/H2B dimers (Luger et al., 1999). For smFRET, H2A-K119C and H2A.Z-K120C histones were labeled with Cy3B-maleimide (Cytiva) prior to octamer and dimer reconstitution, respectively, as described previously (Zhou and Narlikar, 2016). The purified octamers and dimers were diluted 1:1 with freeze buffer (10 mM Tris-HCl, pH 7.4, 2 M NaCl, 40% glycerol, 1mM DTT), flash frozen in aliquots, and stored at −80°C for nucleosome reconstitution and downstream assays. DNA fragments containing an end- or center-positioned 601 nucleosome positioning sequence were generated via Taq PCR amplification in ThermoPol Buffer and purified using a DNA Clean & Concentrator kit (Zymo) as described previously (Singh et al., 2019). Labeled DNA were prepared with 5’-modified primers (IDT) for the following experiments: FAM/Cy3 or FAM/Cy5 for DNA-histone mapping on the 77N4 or 40N40/4N77 template, respectively, biotin-TEG/ATTO 647N for smFRET, and ATTO 647N for fluorescence polarization. Mononucleosomes were reconstituted at 300-1500 nM concentration by salt gradient dialysis as previously described (Luger et al., 1999; Singh et al., 2019). For DNA-histone mapping, nucleosomes were reconstituted without reducing agent in the dialysis buffer using octamers with no cysteine except at the desired crosslinking site.

### Purification of Yeast Chromatin Remodelers

SWR1C and ISW2 were purified from a FLAG-tagged Swr1 or Isw2 yeast strain, respectively, similar to methods described previously (Mizuguchi et al., 2012; Singh et al., 2019) with further modifications: The harvested yeast pellet was pushed through a 60 mL syringe into liquid nitrogen to generate fine frozen noodles. The noodles were gently crushed with pestle and fully lysed using a PM 100 cryomill (Retsch) with 7 x 1 min cycles at 400 rpm. Following the final B-0.1 buffer wash (25 mM HEPES-KOH, pH 7.6, 100 mM KCl, 1 mM EDTA, 2 mM MgCl_2_, 10 mM β-glycerophosphate, 1 mM Na-butyrate, 0.5 mM NaF, 10% glycerol, 0.05% Tween-20, 1mM DTT, protease inhibitors (PIs): 0.05 μM aprotinin, 1 mM benzamidine, 3 μM chymostatin, 4 μM leupeptin, 3 μM pepstatin A, 1 mM PMSF), the remodeler-bound resin was resuspended in B-0.1, transferred into 1.5 mL Eppendorf tubes, centrifuged at 2,500 rpm for 4 min at 4°C, and aspirated. The resin was incubated with 1 mL of 0.5 mg/mL 3x FLAG peptide (Sigma-Aldrich) in B-0.1 and rotated at 4°C for 1 h to elute the tagged remodeler. The elution was collected by centrifugation and repeated. The combined elution was concentrated with a 50 kDa cutoff Amicon Ultra-0.5 mL centrifugal filter (Millipore) by spinning at 14,000 g at 4°C, and then flash frozen in aliquots and stored at -80°C.

Chd1 and RSC were purified using tandem affinity purification (TAP) from a TAP-tagged Chd1 or Rsc2 yeast strain, respectively (Rigaut et al., 1999). 6L of the tagged strains were grown in 2% glucose YPD media to an OD of ∼4 before harvesting by centrifugation at 3,000 rpm for 15 min at 4°C. The pellet was washed by resuspension and centrifugation with cold water, and then with E buffer (20 mM HEPES-NaOH, pH 7.4, 350 mM NaCl, 10% glycerol, 0.1% Tween-20, 1 mM DTT, PIs). The washed pellet was passed through a 60mL syringe into liquid nitrogen to generate frozen noodles, which were lysed by cryomilling. The powder lysate was dissolved in an equal volume of E buffer and clarified by ultracentrifugation at 35,000 rpm for 2 h at 4°C. The supernatant was incubated on a nutator with 300 μL IgG resin (Cytiva) and fresh PIs added for 2.5h at 4°C. The slurry was centrifuged at 700 g for 4 min at 4°C and aspirated. The resin was transferred to an Econo-Pac column (Bio-Rad) and washed three times with 5 mL E buffer without PIs. 300 units TEV protease (in-house prep) in 5 mL E buffer without PIs was added to the resin. The column was capped, wrapped with parafilm, and allowed to nutate overnight at 4°C. The following day, the supernatant was eluted from the column with fresh PIs added. CaCl_2_ was added to a final concentration of 2 mM and the supernatant was added to a disposable column containing 400 μL calmodulin resin (Agilent) pre-equilibrated with E buffer and 2 mM CaCl_2_. The column was capped, parafilm-wrapped, and nutated for 2 h at 4°C. The supernatant was allowed to flow through, and the resin was washed twice with 10 mL E buffer. The resin was incubated in 3 mL E buffer with 10 mM EGTA on the nutator for 10 min at 4°C to elute the tagged remodeler. The elution was concentrated with a 50 kDa cutoff Amicon Ultra 0.5 mL filter via centrifugation at 14,000 g at 4°C. The concentrated elution was dialyzed using a 10 kDa cutoff Slide-A-Lyzer MINI dialysis vessel (ThermoFisher) for 3 h at 4°C into E buffer with 1 mM PMSF and, for RSC only, 50 μM ZnCl_2_. The dialyzed sample was flash frozen in aliquots and stored at -80°C.

All remodeler concentrations were determined via SDS-PAGE using a BSA (NEB) standard titration, followed by SYPRO Ruby (ThermoFisher) staining and quantification using ImageQuant 1D gel analysis.

### Purification of Recombinant Histone Chaperone Chz1

Chz1 was purified via nickel affinity chromatography as follows: a pQE80L plasmid containing Chz1 harboring a N-terminus hexahistidine tag was transformed into E. coli Rosetta 2(DE3)pLysS cells. 3L of transformed cells were grown in 2xYT media to OD 0.7 and induced with 0.8 mM IPTG overnight at 18°C. The cells were harvested the next day by centrifugation at 3,000 rpm for 15 min at 4°C. The pellets were resuspended in 50 mL wash buffer (20 mM Tris-HCl, pH 8.0, 500 mM NaCl, 5 mM BME, 1 mM PMSF) with 10 mM imidazole, sonicated, and clarified by centrifugation at 14,000 rpm for 20 min at 4°C. The supernatant incubated with 150 μL Ni-NTA affinity resin (QIAGEN) on the nutator at 4°C for 3 h. The supernatant was allowed to flow through, and the resin was washed with 5 mL wash buffer containing 10 mM imidazole, 5 mL wash buffer with 40 mM imidazole, and eluted with three rounds of 1.5 mL wash buffer with 200 mM imidazole. The fractions were checked via SDS-PAGE and the cleanest Chz1-containing fraction was concentrated using a 10 kDa cutoff Vivaspin 6 concentrator (Sartorius) at 3,000 g at 4°C. The concentrated Chz1 was dialyzed into storage buffer (20 mM HEPES-NaOH, pH 7.5, 150 mM NaCl, 1 mM TCEP), flash frozen in aliquots, and stored at -80°C. Chz1 concentration was determined to be ∼100 μM by UV absorbance at 276 nm.

### Site-Directed DNA-Histone Mapping

Site-directed crosslinking to map DNA-histone contacts was performed as previously described (Kassabov et al., 2003; Nodelman et al., 2017). 1-1.5 μM H2B-Q59C or H2B-S93C nucleosomes with FAM/Cy3- or FAM/Cy5-conjugated DNA were labeled with 200-400 μM 4-azidophenacyl bromide (APB) prepared fresh as a 80mM stock dissolved in N,N-dimethylformamide (DMF). The nucleosomes were labeled for 3 h in the dark at room temperature before being quenched with 5 mM DTT and stored on ice. The crosslinking reactions were prepared in 50 μL volume with 100-150 nM APB-labeled nucleosomes, 300 nM remodeler, 450 nM H2A.Z-H2B dimer for the SWR1C experiments, and 1 mM nucleotide in reaction buffer (For SWR1C: 25 mM HEPES-KOH, pH 7.6, 70 mM KCl, 0.2 mM EDTA, 5 mM MgCl_2_, 5% glycerol, 1 mM DTT, 0.1 mg/mL BSA; for Chd1/ISW2/RSC: 20 mM Tris-HCl, pH 7.5, 50 mM KCl, 5 mM MgCl_2_, 5% sucrose, 1 mM DTT, 0.1 mg/mL BSA). ADP stock was prepared by incubating a 100 mM stock to a final concentration of 44 mM with 1 M (18 %) glucose, 0.1 U/ul hexokinase, and 5 mM MgCl_2_ for 20 min at room temperature prior to use. ADP•BeF_3_^-^ was prepared by adding to a final concentration of 1 mM ADP, 6 mM NaF, 1.2 mM BeCl_2_, and 2.5 mM MgCl_2_ to the reaction buffer. The reactions were incubated at room temperature for 15-30 min. The reactions were transferred to a 96-well UV-transparent plate (Corning) and irradiated with a UV TransIlluminator (VWR) at 302 nm for 15 sec to crosslink. The reactions were transferred back into Eppendorf tubes and mixed with 100 μL quench buffer (reaction buffer with 5 mM EDTA, 5 mM DTT) and 150 μL post-irradiation buffer (20 mM Tris-HCl, pH 8, 50 mM NaCl, 0.2 % SDS). The SWR1C dimer exchange reaction was initiated by adding 3 μM ATP and immediately UV-crosslinked and mixed with quench and post-irradiation buffer at the appropriate time points. The samples were vortexed and incubated for 20 min at 70 °C. The incubated samples were added 300 μL 5:1 phenol:chloroform (Sigma-Aldrich), vortexed, and centrifuged for 2 min at 16,100 g at room temperature. ∼250 μL of the top aqueous layer was removed from each sample without disturbing the aqueous-organic interface and 280 μL of wash buffer (1 M Tris-HCl, pH 8, 1 % SDS) was added to the sample, which was then vortexed and centrifuged. This wash was repeated three more times. Crosslinked DNA was precipitated by adding 1.5 μL 10 mg/mL sonicated salmon sperm DNA (Agilent), 33 μL 3 M sodium acetate, pH 5.2, and 750 μL 100% EtOH. The samples were vortexed and incubated on ice at 4 °C overnight. The next day, the precipitated DNA was pelleted at 16,100 g for 30 min at 4°C. The supernatant was carefully removed with a pipet. The pellet was washed with 750 μL 70% EtOH and centrifuged at 16,100 g for 5 min at 4°C twice. The pellet was air-dried by inverting the opened Eppendorf tube for at least 1 h in the dark. The dried pellet was resuspended with 100 μL resuspension buffer (20 mM ammonium acetate, 0.1 mM EDTA, 2% SDS. The sample was vortexed for 30 s and centrifuged at 16,100 g for 10 min at room temperature. The supernatant was transferred to another Eppendorf tube and incubated for 2 min at 90°C. The sample was pulsed, added 5 μL 2 M NaOH, vortexed, and incubate for 45 min at 90°C to cleave the crosslinked DNA. The sample was pulsed to collect any condensate and added 105 μL 20 mM Tris-HCl, pH 8 and 6 μL 2 M HCl. The sample was then vortexed, added 2 μL 1 M MgCl_2_ and 480 μL 100% EtOH, vortexed again, and left to precipitate overnight at -20°C. The following day, the sample was pelleted at 16,100 g for 30 min at 4°C. The supernatant was carefully removed with a pipet. The pellet was washed with 750 μL 70% EtOH and centrifuged for at 16,100 g for 5 min at 4°C twice. The pellet was air-dried by inverting the opened Eppendorf tube for at least 1 h in the dark. The dried pellet was resuspended with 4 μl of 90% formamide, 10 mM NaOH, 1 mM EDTA, and 0.01% bromophenol blue. The sample was vortexed and incubate for 3 min at 90°C. The heated sample was pulsed and cooled at room temp for 1 min before being loaded onto a denaturing 8% polyacrylamide sequencing gel. A G+A sequencing ladder was used as reference to identify the DNA crosslink location. The sequencing ladder was prepared as previously described (Maxam and Gilbert, 1980) from the corresponding fluorescently labeled DNA used to reconstitute the nucleosome. The gel was run at 65 W for 1.5 h and visualized on a Typhoon Imager by scanning at 473 nm (FAM), 532 nm (Cy3), or 635 nm (Cy5). The dimer exchange crosslinking time course was quantified using ImageQuant 1D gel analysis by normalizing crosslinked band to the uncut DNA band for each time point. Full gel images are provided in Supplementary Figure S2.

### smFRET Imaging and Analyses

#### Flow Cell Preparation

Glass coverslips were placed in coplin jars and cleaned by sonicating for 30 minutes in methanol After washing with copious amounts of DI H_2_O, piranha solution (3:1 mixture of sulfuric acid and 30% hydrogen peroxide) was added to the coplin jars and allowed to sit for 1 hour. The piranha solution was removed, and coverslips were again washed with DI H_2_O. The coverslips were then functionalized with a mixture of methoxypolyethylene glycolsuccinimidyl valerate, MW 5,000 (mPEG-SVA-5000; Laysan Bio, Inc.) and biotin-methoxypolyethylene glycol-succinimidyl valerate, MW 5,000 (biotin-PEG-SVA-5000; Laysan Bio, Inc.) as previously described (Graham et al., 2017).

Microfluidic chambers were assembled as follows: a diamond-tipped rotary bit was used to drill holes 10 mm apart in a glass microscope slide; a 4.5 mm wide piece of double-sided SecureSeal Adhesive Sheet (Grace Bio-Labs) was placed parallel to one side of holes and across the slide, a second 4.5 mm wide piece of double-sided placed on the other side of the holes making a channel, a piece of functionalized coverslip was then secured to the second side of the adhesive sheet and the edges of the coverslip were sealed with epoxy (Devcon). To make the microfluidic chamber PE20 tubing was inserted into one hole and PE60 tubing into the other (Intramedic), and the tubing was fixed in place with epoxy. The microfluidic cells were stored under vacuum until time of use.

#### Preparation of Calibration DNA

The calibration substrate used for channel alignment consists of Cy5 and Cy3 labeled 60 bp duplex. The substrate was made by mixing 10 μM of oligos (IDT) oAM200, oTG415, and oTG416 (Supplementary Table S4) in 100 μL of 20 mM Tris, 300 mM NaCl, 1 mM EDTA pH 8.0. The sample was then placed in a 2-liter beaker containing water heated to 90°C and allowed to cool overnight to room temperature.

#### smFRET Imaging

Imaging of single-molecule nucleosomes was achieved using a through-objective TRIF microscope configured around an inverted Olympus IX-71 microscope. Each laser beam 532 nm (Coherent Sapphire 532) and 641 nm (Cube 641) were first expanded and then combined using dichroic mirrors. The combined beams were expanded again and focused onto the rear focal plane of an oil-immersion objective (Olympus UPlanSApo, 100 3; NA, 1.40). To achieve TRIF illumination the focusing lens was manually translated in the vertical plane. A multipass dichroic mirror was used to separate emission from excitation light. The emission light was then sent across a StopLine 488/532/635 notch filter (Semrock) to further reduce excitation light. A home-built beamsplitter (Graham et al., 2017) was used to separate emission from Cy3B and ATTO 647N and the then imaged on separate halves of EMCCD camera (Hamamatsu, ImageEM 9100-13) operating at maximum EM gain. The focus was adjusted manually, and the sample was positioned on the microscope using an automated microstage (Mad City Labs).

For calibration data the flow cell chamber was incubated with 35 μL 0.71 mg/mL streptavidin (25 μL 1 mg/mL streptavidin in 1x PBS diluted with 10 μL of 10 mM Tris-HCl, pH 7.4, 1 M NaCl) for 5 min. The calibration substrate, 5-prime biotinylated 60 bp dsDNA labeled with Cy5 and Cy3, was diluted to ∼30 pM in Tris pH 7.4 (10 mM), NaCl (1M), EDTA (1 mM), protocatechuic acid (5 mM), protocatechuate 3,4-dioxygenase (0.1 mM), and Trolox (1 mM). The biotinylated, fluorescent DNA substrate was immobilized on a glass coverslip in a microfluidic chamber. Images were acquired of different fields of view (∼120) with 0.5 sec simultaneous exposure to 532 and 641 nm lasers using a surface power density of 4 mW/cm^2^ for the 532 nm laser and 2.4 mW/cm^2^ for the 641 nm laser.

The smFRET data was collected at a surface power density of 0.6 mW/cm^2^ for 532 nm and 0.4 mW/cm^2^ for 641nm. The integration time was 0.5 sec per frame and the images were collected continuously at a cycle of 4 frames of 532 nm excitation and 1 frame of 641 nm. The flow cell chamber was incubated with 35 μL 0.71 mg/mL streptavidin (25 μL 1 mg/mL streptavidin in 1x PBS diluted with 10 μL of 10 mM Tris-HCl, pH 7.4, 1 M NaCl) for 15 min. The chamber was passivated with 150 μL smFRET wash buffer (25 mM HEPES-KOH, pH 7.6, 70 mM KCl, 0.2 mM EDTA, 5 mM MgCl_2_, 1 mM DTT, 0.2 mg/mL acetylated BSA (Promega), 0.02% NP-40 (Sigma-Aldrich)) for 5 min. The chamber was then equilibrated with 150 μL smFRET reaction buffer (20 mM HEPES-KOH, pH 7.6, 56 mM KCl, 0.16 mM EDTA, 4 mM MgCl_2_, 10% glycerol, 1 mM DTT, 0.1 mg/mL acetylated BSA, 0.02% NP-40, 1mM Trolox, 0.8% glucose, 0.24 mg/mL glucose oxidase, 0.24 mg/mL catalase acetylated BSA 0.2 mg/mL, 0.02% NP-40). For dimer eviction reactions, 50 μL of 90 pM biotinTEG-117N4-ATTO 647N, Cy3B-labeled H2A nucleosome, 810 pM biotinTEG-117N0 nucleosome, 10 nM 77N0 nucleosome, 30 nM SWR1C, 70 nM H2A.Z-H2B dimer, and 70 nM Chz1 in reaction buffer was flowed into the chamber and incubated for 5 min. The chamber was then washed with 150 μL wash buffer. The dimer eviction reaction was initiated 20 sec after beginning imaging by injecting 50 μL reaction buffer containing the indicated concentration of ATP, 10 nM 77N0 nucleosome, 30 nM SWR1C, 70 nM H2A.Z-H2B dimer, and 70 nM Chz1. The imaging data was collected for a total of 6 min. For dimer deposition reactions, 50 μL of 90 pM biotinTEG-117N4-ATTO 647N nucleosome, 810 pM biotinTEG-117N0 nucleosome, 5 nM 77N0 nucleosome, 25 nM SWR1C, 50 nM Cy3B-labeled H2A.Z-H2B dimer, and 50 nM Chz1 in reaction buffer was first incubated in the chamber and the deposition reaction was initiated with 50 μL reaction buffer containing 200 μM ATP, 10 nM 77N0 nucleosome, 25 nM SWR1C, 50 nM Cy3B-labeled H2A.Z-H2B dimer, and 50 nM Chz1. All smFRET experiments were performed at room temperature.

#### Alignment of the Donor and Acceptor Channels

Using custom MATLAB scripts, an automated spot-detection algorithm was used to identify fluorescent molecules in the 532 and 641 emission channels of each image. Spots were identified by first subtracting local background and identifying particles based on local maximum. For greater precision in location, shape, and amplitude, particles were fit using a 2D Gaussian. For each image spots whose coordinates < 6 pixels from surrounding molecules and are within 6 pixels of a spot in the corresponding channel are added to the initial calibration list. The initial calibration list is then refined based on spot diameter and amplitude. A final calibration list is made using the method outlined in the paper from (Friedman and Gelles, 2015). Briefly, for each image the coordinates (x_1_,y_1_) of a spot in channel 1 (532 emission) is mapped onto coordinates (x_2_,y_2_) in channel 2 (641 emission) to identify a matching partner spot using the transformation Eq (1)

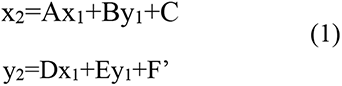

where A–F are fit parameters. Because of systematic various across the field of view, the fit parameters are determined for each spot coordinate (x_1_, y_1_) by fitting pairs of corresponding points from the initial calibration list that are within a 30 x 30-pixel box of (x_1_, y_1_). A spot that maps greater than 2 pixels from all spots in channel 2 is removed form calibration list. This process is repeated for spots in channel 2, mapping coordinates onto channel 1 using the updated calibration list. Channel 2 spot that are within 1 pixel of to its channel 1 partner are retained in the calibration list. The calibration list is further refined by repeating the above process removing spots that are greater than 1 pixel from spots in the other channel to give the final calibration list that is used to map coordinates between the two channels.

#### Detection and Quantification of Eviction Events

Quantification of single-molecule eviction experiments were performed using the following steps. Spot detection was achieved using an image of the 532 nm channel and 641 nm channel from the start of the movie with the method described above. Molecules within a radius of 6 pixels of other particles were excluded from the analysis to avoid crosstalk. Fluorescent pairs were identified using the calibration list and transformation method described above. Stage drift was estimated in movies using average change in x and y position between successive frames for particles in the 641 nm channel and region of interest (ROI) positions were translated to compensate. Integrated intensities derived from raw images were used for energy transfer efficiency calculations. The integrated intensity of each particle over time was calculated as the summation of a circular ROI of radius 4 pixels centered on the particle. The local background intensity was determined as the median intensity for pixels flanking the ROI. Photobleaching events for ATTO 647N were automatically identified using the built-in MATLAB function findchangepts, which identifies abrupt changes in signal, to first mark changes and aggregate intensities past the change point for all molecules. Next, the aggregated intensity histogram was fit to a Gaussian distribution, and a threshold for ATTO 647N photobleaching was set at two standard deviations above the mean. A similar method was used to define photobleaching threshold (either due to photobleaching or loss of Cy3B by eviction from the nucleosome) for the total Cy3B emission signal, which is the sum of intensity for Cy3B emission and Cy3B excited ATTO 647N emission. The trajectories for each molecule were then automatically truncated based on photobleaching events and if necessary, the point of truncation was manually adjusted. For each ATP condition the trajectories from a minimum of 3 independent experiments were combined. The initial energy transfer for each molecule was defined as the median transfer efficiency of the first 10 frames. The trajectories with an initial energy transfer greater than 0.4 were globally fitted to a three-state Hidden Markov Model using the ebFRET MATLAB software package (van de Meent et al., 2014) with default parameters. A custom MATLAB script was used to identify eviction events, the duration of the priming, eviction, and release intervals based on the output FRET states from ebFRET. Proximal first eviction events were defined as change a from an initial E.T. state between 0.58 and 0.77 to a E.T. state between 0.4 and 0.57. Distal second eviction events were defined when following a proximal eviction there is a change a from an E.T. state between 0.4 and 0.57 to a E.T. state below 0.35 without loss of Cy3B signal. Distal first eviction events were defined as change a from an initial E.T. state between 0.58 and 0.77 to an E.T. state above 0.77. Proximal second eviction events were defined when following a Distal eviction there is a change a from an E.T. state above 0.78 to an E.T. state below 0.35 without loss of Cy3B signal. Single-High eviction events were defined as change a from an initial E.T. state above 0.77 to an E.T. state below 0.36 without loss of Cy3B signal. Single-Middle eviction events were defined as change a from an initial E.T. state between 0.58 and 0.77 to an E.T. state below 0.35 without loss of Cy3B signal. Single-Low eviction events were defined as change a from an initial E.T. state between 0.4 and 0.57 to an E.T. state below 0.35 without loss of Cy3B signal. The length of time for eviction was determined by first using a 3-point moving median to smooth the E.T. signal and next fitting a baseline to the initial E.T. state and the E.T. state following eviction. The start and stop of the eviction were defined as the point of 5% deviation of smooth E.T. signal from respective baselines. The duration of the priming interval is then defined as the time from the start of the trajectory to the start of eviction. For eviction events that result in a E.T. state below 0.35, the release interval was demarcated as the length of time between the end of eviction to the loss of Cy3B signal.

For each type of eviction event the distribution in the duration of priming interval were fit to a single exponential using the expfit function in MATLAB. To calculate the half-life of the release phase for each ATP concentration, survival curves of the time for release (Supplementary Figure S3) were constructed using the Kaplan-Meier procedure. Release intervals were classified as right-censored if no donor loss event was observed before the end of the observation interval. Statistical analysis of the priming, eviction, and release interval between the different ATP concentrations (0.5, 5, 100 μM ATP) were performed using an ordinary ANOVA test in Prism Version 9.3.1, P values % 0.05 were considered significant for this analysis. Because of the shape of the priming and release time distributions, the statistical tests were performed on the logarithm base 2 scale to better satisfy the assumption of homoscedasticity. To obtain the kymographs shown in Figure 2C, energy transfer efficiency trajectories were truncated upon photobleaching and then pooled. Survival curves of the energy transfer (Figure 2B) were constructed using the Kaplan-Meier procedure. Trajectories were classified as right-censored if the molecule remained in the high E.T. state at the end of the observation interval.

To examine unwrapping events that occurred during the priming interval, trajectories were truncated at the start of eviction. Molecules with an initial energy transfer greater than 0.6 were globally fitted to a three-state Hidden Markov Model using the ebFRET MATLAB software package (van de Meent et al., 2014) with default parameters. A custom MATLAB script was used to identify unwrapping events and the duration of the events. Unwrapping events were defined as a transition from an initial state with a median E.T. greater than 0.6 to state with a median E.T. less than 0.45 and a subsequent return to a state with a median E.T. greater than 0.6. Transition density plots were made by first normalizing the intensities of transition to the total number of frames, where intensities indicate the number of transitions between E.T. X and Y per unit time. Intensities that are greater than 0.0005 were plotted using the contour function in MATLAB. Statistical analysis between (-) ATP and (+) ATP (100 μM ATP) were performed using an unpaired t test in Prism Version 9.3.1, P values % 0.05 were considered significant for this analysis.

#### Detection and Quantification of Deposition Events

Analysis of single-molecule deposition of Cy3B-labeled H2A.Z experiments were performed using the following steps. The ROIs for the tethered nucleosomes were determined with an image of the 641 nm channel from the start of the movie using the method described in *Alignment of the Donor and Acceptor Channels.* To examine the non-specific interaction between H2A.Z and the coverslip, control ROIs were picked that were > 6 pixels from tethered nucleosomes. The position of ROIs in the Cy3B channel for tethered nucleosomes and control ROIs were identified using the calibration list and transformation method described in *Alignment of the Donor and Acceptor Channels*. Stage drift was estimated using average change in x and y position between successive frames for particles in the 641 nm channel and ROI positions were translated to compensate. In each of the 532 nm excitation frames, Cy3B-labeled H2A.Z localized spots were detected using the algorithm described in *Alignment of the Donor and Acceptor Channels* with the following modification. The background peak of the Cy3B intensity histogram was fit locally to a Gaussian distribution and a background threshold was calculated as one standard deviation above the mean. Localized H2A.Z spots with an intensity above background threshold were fit to a 2D Gaussian. Integrated intensities of the total Cy3B emission signal (sum of intensity for Cy3B emission and Cy3B excited ATTO 647N emission) was used to identify binding and dissociation of H2A.Z to tethered nucleosomes using the following method. The trajectories for Cy3B were first denoised using a piecewise constant approximation (Little and Jones, 2011) followed by identification of changepoints using the built-in MATLAB function findchangepts. Binding events were required to satisfy the following conditions: (1) the intensity must increase by 30% at identified changepoint, (2) the intensity must be two standard deviations above the mean Cy3B background signal, (3) the localized H2A.Z must be within 1.25 pixels of the center of the ROI. The dissociation events were defined as changepoints with a greater than 50% decrease in amplitude, which is less than one standard deviations above the mean Cy3B background signal. Trajectories were manually annotated to mark deposition events identified by an anticorrelated increase in Cy3B excited ATTO 647N emission and a decrease in Cy3B emission. To obtain the fraction bound plot (**Figure S2**), the initial binding events are sorted by arrival time and the cumulative sum of these events are normalized to the number of surface-tethered nucleosome complexes. The dwelltime distribution for H2A.Z colocalized with surface-tethered nucleosomes was fit to a double exponential using the MEMLET MATLAB software package (Woody et al., 2016). A bootstrap analysis (1000 iterations) was used to estimate the confidence intervals for the exponential fit.

### Fluorescence Polarization Nucleosome Binding Assay

The fluorescence polarization binding assay was performed as follows: A 2-fold serial dilution of SWR1C-nucleosome binding reactions was prepared in reaction buffer (25 mM HEPES-KOH pH 7.6, 70 mM KCl, 0.2 mM EDTA, 5 mM MgCl_2_, 1 mM DTT, 0.1 mg/mL BSA) to a final concentration of 0-250 nM SWR1C, 10 nM 77N4-ATTO 647N nucleosome, 1 mM nucleotide in 20 μL reaction volume. The reactions were transferred onto a 384-well black microplate (PerkinElmer) and incubated for 20 minutes at room temperature. The fluorescence polarization signal was measured in a Tecan Spark microplate reader using an excitation wavelength of 631 nm and emission wavelength of 686 nm. A binding curve was generated from the SWR1C titration, normalized, and fitted to the quadratic binding equation

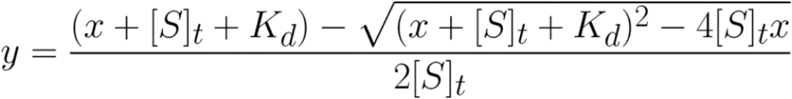

where [S]_t_ is the nucleosome concentration (10 nM), x is the SWR1C concentration, y is the fraction of nucleosome bound, and K_d_ the dissociation constant to be determined from the fit. At least three replicates were generated for each condition and a global fit on the replicates were performed using OriginLab to calculate the K_d_.

## Supplemental Figures

**Figure S1.** Site-directed DNA-histone mapping on different nucleosome templates. (A) Chd1 binding in apo or ADP-bound state induces a 2-nt shift at SHL5.5 on a 40N40 nucleosome. (B) RSC does not exhibit any translocation in its apo or ADP-bound form, but does translocate with a 10bp periodicity in the presence of 1 mM AMP-PNP. (C) SWR1C does not translocate DNA on the opposite SHL5 of the 40N40 nucleosome relative to the side shown in Figure 1A. (D) SWR1C does not alter the DNA path near its predicted binding site on the linker-distal side of a 4N77 nucleosome. (E) SWR1C does not change the path of nucleosomal DNA on the linker-proximal side of an asymmetric nucleosome regardless of 601 positioning sequence asymmetry. (F) SWR1C does not induce any translocation at SHL3 using an alternative crosslinking site at H2B residue 93, unlike ISW2. (G) SWR1C does not induce any translocation at SHL3.5 of the guide strand complement to the template strand that SWR1C is predicted to bind. (H) DNA-histone mapping detecting SWR1C dimer exchange on the linker-proximal side of an asymmetric nucleosome.

**Figure S2.** Full sequencing gel images for DNA-histone mapping in Figures 1 and S1. (A) Extended gel of Figures 1B left panel (left) and S1C left panel (right) scanning at 473 nm and 635 nm, respectively. (B) Extended gel of Figure 1B middle panel scanning at 473 nm. (C) Extended gel of Figures 1B right panel and S1B (left) and S1C right panel (right) scanning at 473 nm and 635 nm, respectively. (D) Extended gel of Figure S1A scanning at 473 nm. (E) Extended gel of Figures 1C (left) and S1E left panel (right) scanning at 473 nm and 532 nm, respectively. (F) Extended gel of Figures 1D (left) and S1H (right) scanning at 473 nm and 635 nm, respectively. (G) Extended gel of Figures S1D (left gel; left half) and S1F left two panels (left gel; right half) scanning at 473 nm and Figures S1E right panel (right gel; left half) and S1G left two panels (right gel; right half) scanning at 635 nm. The “G+A” lanes each indicate a G+A sequencing ladder generated using the corresponding nucleosome DNA template.

**Figure S3.** The number of eviction events and duration of the priming and release phase is dependent on ATP concentration. (A) Observed events for each type of eviction with indicated ATP concentration. (B) Duration of priming phase for each type of eviction with indicated ATP concentration. (C) Duration of release phase for each type of eviction with indicated ATP concentration. (D) Duration of priming and release phase for aggregate eviction events with indicated ATP concentration. (B-D) Duration values derived from fitting distributions to a single exponential and error bars represent 95 % C.I.

**Figure S4.** SWR1C-nucleosome binds and deposits H2A.Z within the nucleosome (A) Schematic of of the smFRET H2A.Z deposition assay. (B) Example trajectory highlighting binding of H2A.Z (*) and deposition (#). The H2A.Z labeled with Cy3B donor fluorophore was excited, and donor emission (green) and nucleosome labeled with ATTO 647N acceptor emission (red) were recorded (top panel) and used to calculate energy transfer efficiency (blue, bottom panel). (B) (right side) Rastergrams summarize colocalization traces of individual SWR1C-nucleosome complexes (N=307) each in a single row and sorted according to the arrival time of H2A.Z. (left side) Rastergrams summarize traces for individual background ROI (N=. The events are colored by dwell time. (C) Fraction of SWR1C-nucleosome complexes that are bound by H2A.Z sorted by time arrival time. (D) Distribution of H2A.Z dwelltime, red line represents the fit to double exponential, and black dashed line corresponds to 95 % C.I. from bootstrap analysis.

**Figure S5.** Sample Single-Molecule colocalization and deposition of H2A.Z with the SWR1C-nucleosome complex, related to Figure S3. Example trajectory highlighting binding of H2A.Z (*) and deposition (#). The H2A.Z labeled Cy3B donor fluorophore was excited, and donor emission (green) and nucleosome labeled ATTO 647N acceptor emission (red) were recorded (top panel) and used to calculate energy transfer efficiency (blue, bottom panel).

