## Supplemental figures and tables for "H2A.Z deposition by SWR1C involves multiple ATP-dependent steps"

### Figure S1

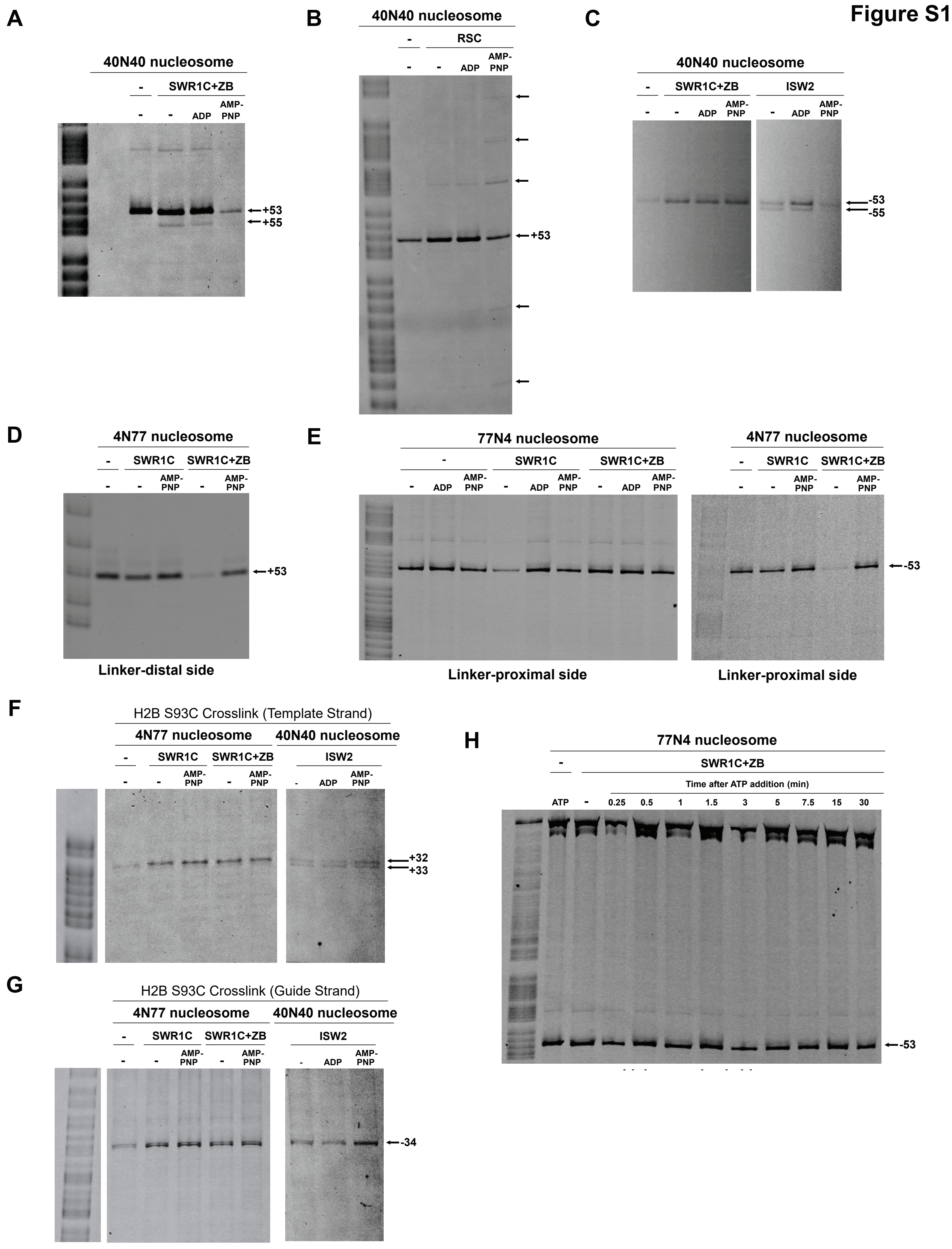
A
40N40 nucleosome
-
SWR1C+ZB
-
-
ADP
AMP-PNP
+53
+55
B
40N40 nucleosome
-
RSC
-
-
ADP
AMP-PNP
+53
C
40N40 nucleosome
-
SWR1C+ZB
ISW2
-
-
ADP
AMP-PNP
-
ADP
AMP-PNP
-53
-55
D
4N77 nucleosome
-
SWR1C
SWR1C+ZB
-
-
AMP-PNP
-
AMP-PNP
+53
Linker-distal side
E
77N4 nucleosome
-
SWR1C
SWR1C+ZB
-
ADP
AMP-PNP
-
ADP
AMP-PNP
-
ADP
AMP-PNP
Linker-proximal side
4N77 nucleosome
-
SWR1C
SWR1C+ZB
-
-
AMP-PNP
-
AMP-PNP
-53
Linker-proximal side
F
H2B S93C Crosslink (Template Strand)
4N77 nucleosome
40N40 nucleosome
-
SWR1C
SWR1C+ZB
ISW2
-
-
AMP-PNP
-
AMP-PNP
-
ADP
AMP-PNP
+32
+33
G
H2B S93C Crosslink (Guide Strand)
4N77 nucleosome
40N40 nucleosome
-
SWR1C
SWR1C+ZB
ISW2
-
-
AMP-PNP
-
AMP-PNP
-
ADP
AMP-PNP
-34
H
77N4 nucleosome
-
SWR1C+ZB
Time after ATP addition (min)
ATP
-
0.25
0.5
1
1.5
3
5
7.5
15
30
-53

**Figure S2**

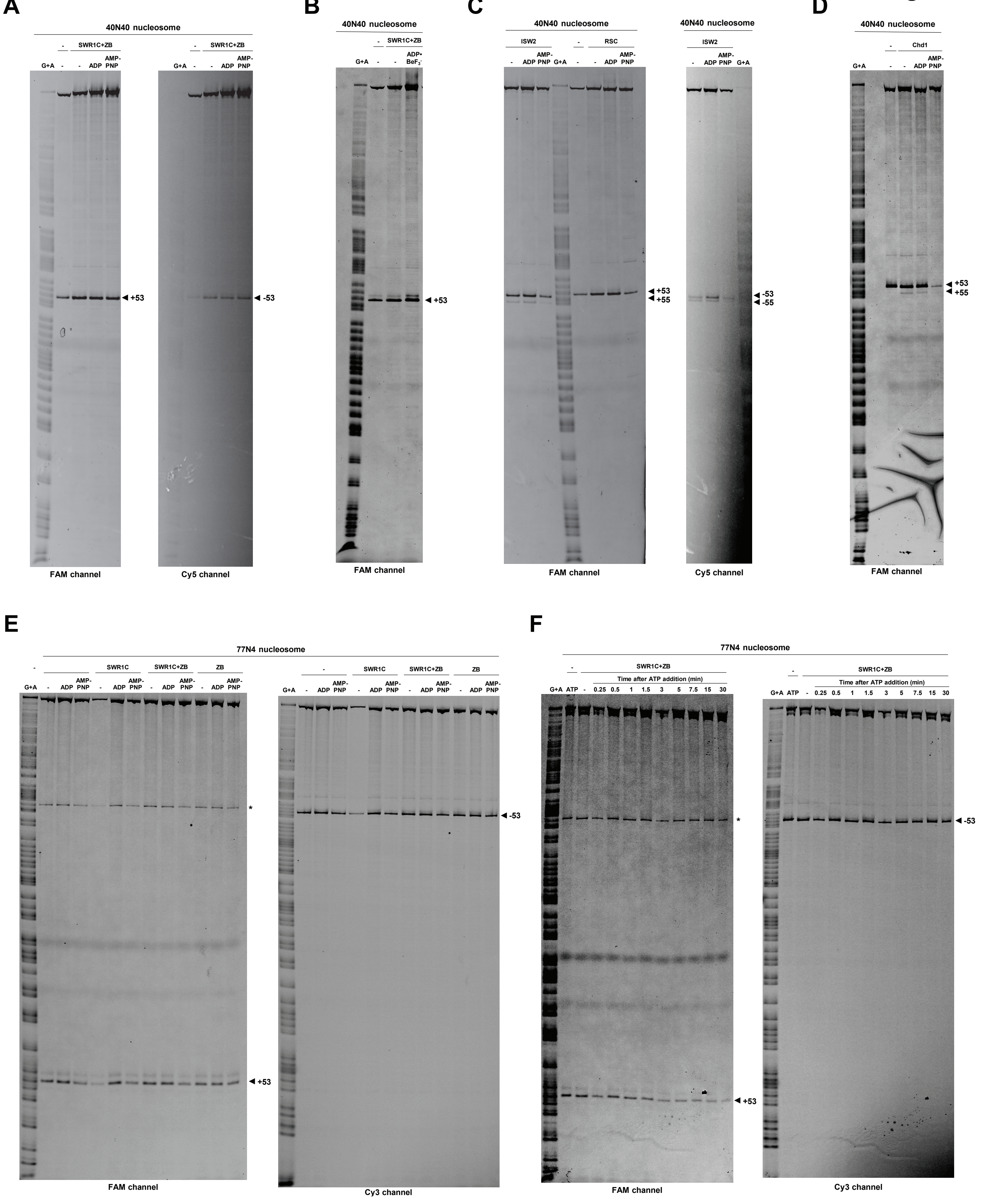
A
40N40 nucleosome
SWR1C+ZB
G+A
-
ADP
AMP-PNP
◄ +53
◄ -53
FAM channel
Cy5 channel
B
40N40 nucleosome
SWR1C+ZB
ADP•BeF3-
◄ +53
FAM channel
C
40N40 nucleosome
ISW2
RSC
◄ +53
◄ +55
◄ -53
◄ -55
FAM channel
Cy5 channel
D
40N40 nucleosome
Chd1
◄ +53
◄ +55
FAM channel
E
77N4 nucleosome
SWR1C
SWR1C+ZB
ZB
*
◄ +53
◄ -53
FAM channel
Cy3 channel
F
77N4 nucleosome
SWR1C+ZB
Time after ATP addition (min)
G+A
ATP
0.25
0.5
1
1.5
3
5
7.5
15
30
*
◄ +53
◄ -53
FAM channel
Cy3 channel

### Figure S2

## G

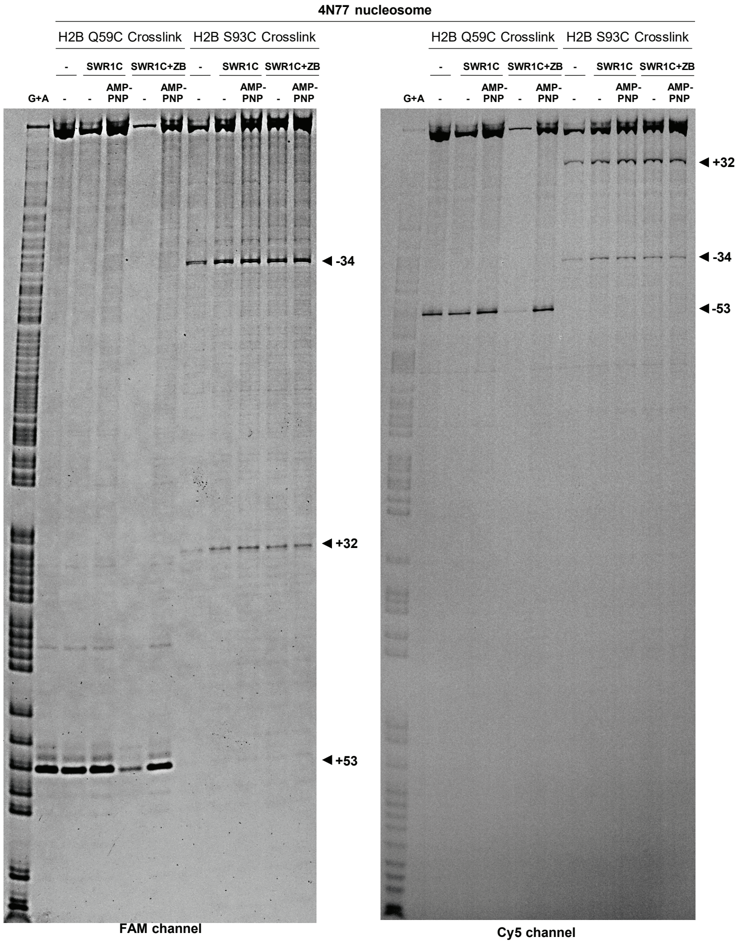

4N77 nucleosome
H2B Q59C Crosslink
H2B S93C Crosslink
SWR1C
SWR1C+ZB
G+A
AMP-PNP
-34
+32
+53
-53
FAM channel
Cy5 channel

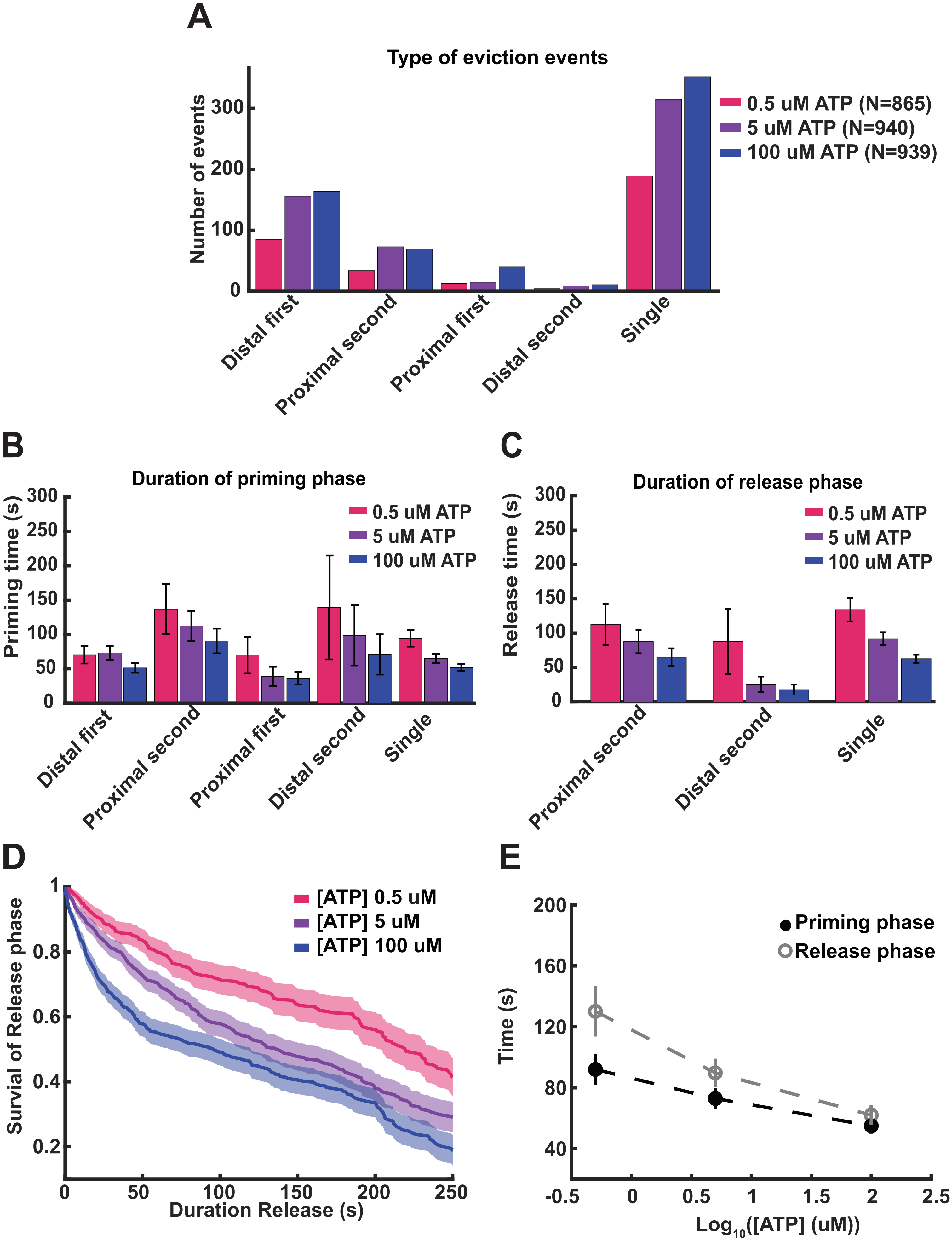
A
Type of eviction events
Number of events
0.5 uM ATP (N=865)
5 uM ATP (N=940)
100 uM ATP (N=939)
Distal first
Proximal second
Proximal first
Distal second
Single
B
Duration of priming phase
Priming time (s)
0.5 uM ATP
5 uM ATP
100 uM ATP
C
Duration of release phase
Release time (s)
D
Survial of Release phase
Duration Release (s)
[ATP] 0.5 uM
[ATP] 5 uM
[ATP] 100 uM
E
Time (s)
Priming phase
Release phase
Log10([ATP] (uM))

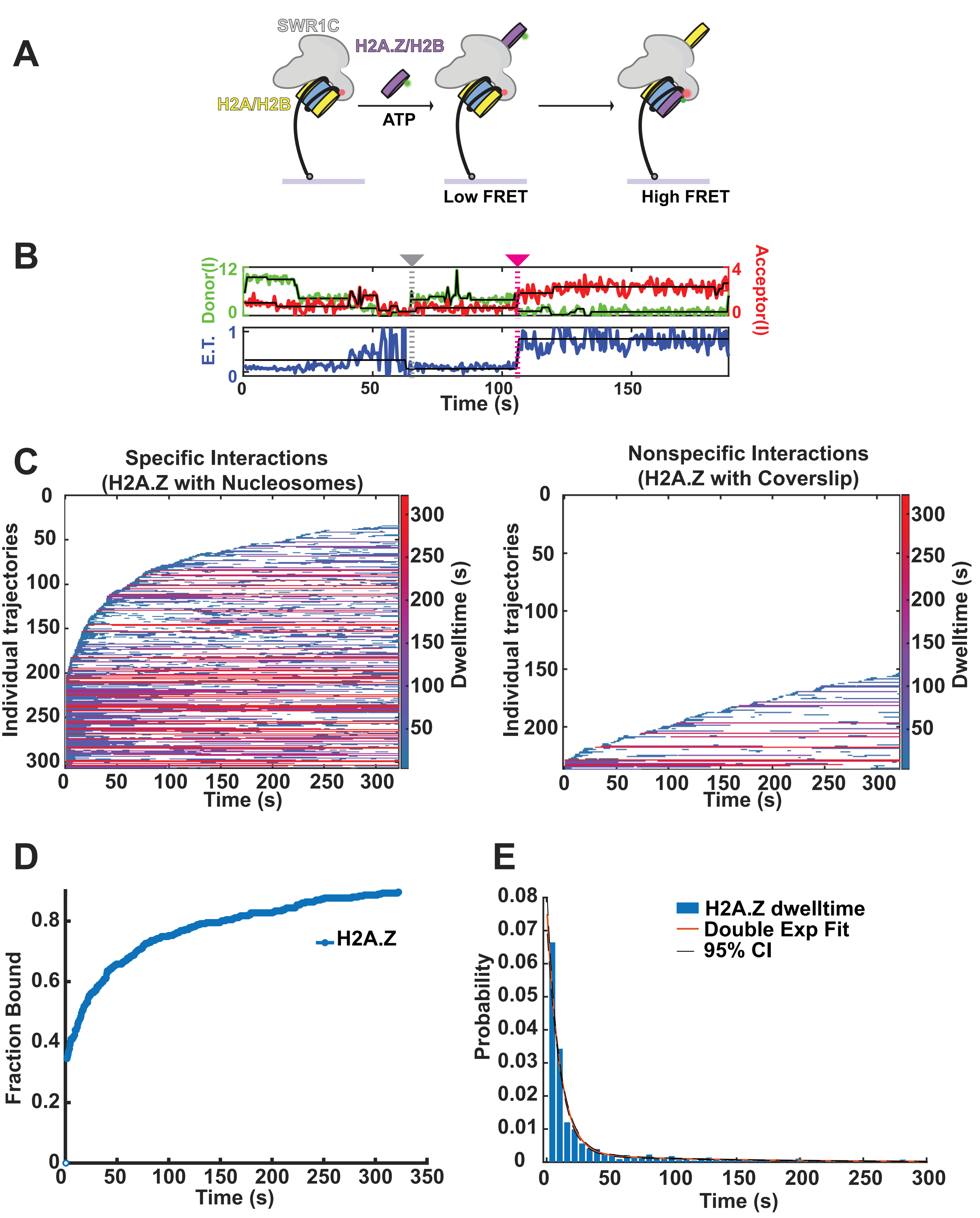
A
SWR1C
H2A.Z/H2B
H2A/H2B
ATP
Low FRET
High FRET
B
Donor(I)
12
0
4
0
Acceptor(I)
E.T.
1
0
0
50
100
150
Time (s)
C
Specific Interactions
(H2A.Z with Nucleosomes)
Individual trajectories
Dwelltime (s)
Time (s)
Nonspecific Interactions
(H2A.Z with Coverslip)
Individual trajectories
Dwelltime (s)
Time (s)
D
Fraction Bound
H2A.Z
Time (s)
E
Probability
H2A.Z dwelltime
Double Exp Fit
95% CI
Time (s)

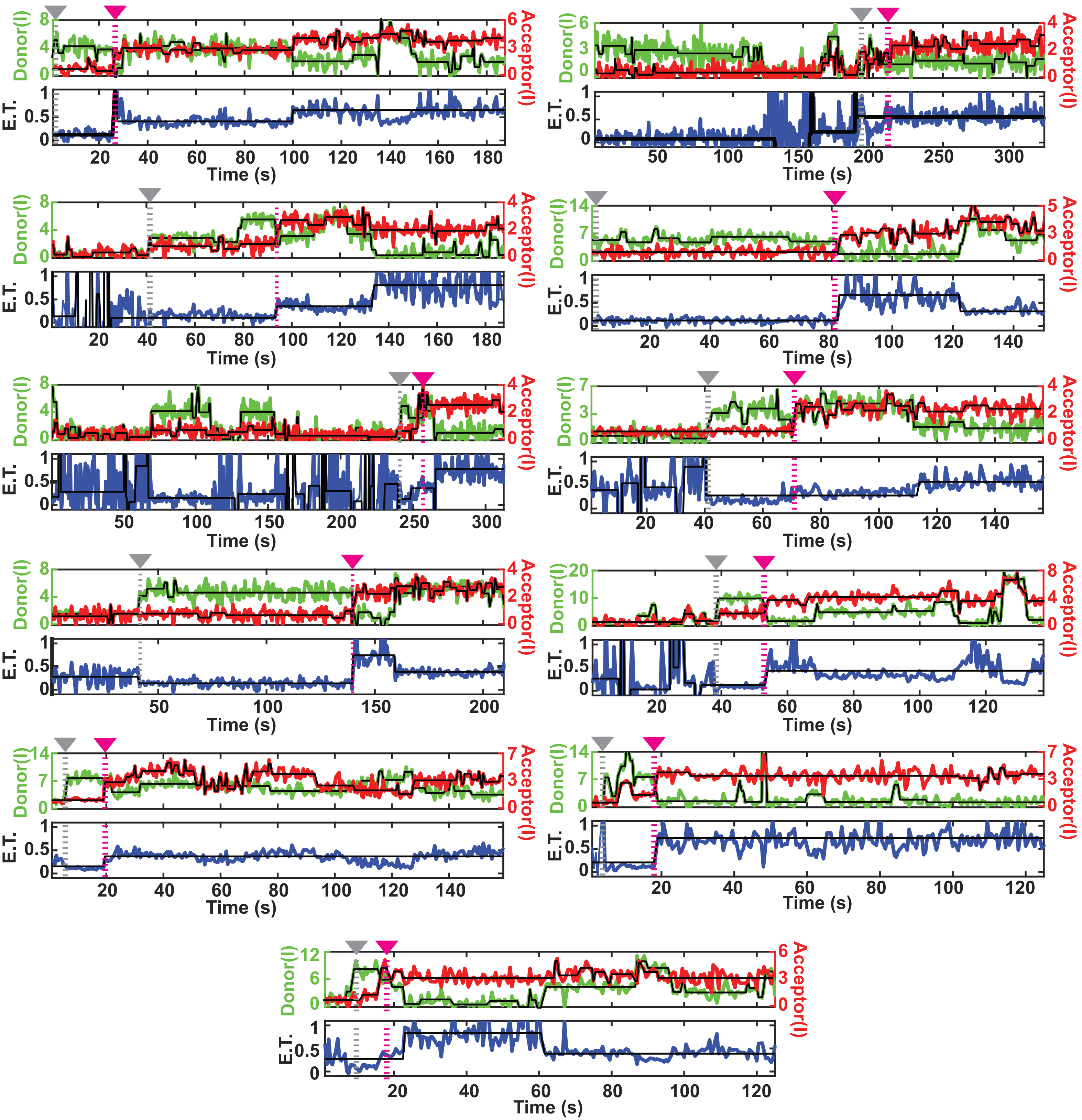

Donor(I)
Acceptor(I)
E.T.
Time (s)

#### Supplemental Figure Legends

**Figure S1**. Site-directed DNA-histone mapping on different nucleosome templates. (A) Chd1 binding in apo or ADP-bound state induces a 2-nt shift at SHL5.5 on a 40N40 nucleosome. (B) RSC does not exhibit any translocation in its apo or ADP-bound form, but does translocate with a 10bp periodicity in the presence of 1 mM AMP-PNP. (C) SWR1C does not translocate DNA on the opposite SHL5 of the 40N40 nucleosome relative to the side shown in Figure 1A. (D) SWR1C does not alter the DNA path near its predicted binding site on the linker-distal side of a 4N77 nucleosome. (E) SWR1C does not change the path of nucleosomal DNA on the linker-proximal side of an asymmetric nucleosome regardless of 601 positioning sequence asymmetry. (F) SWR1C does not induce any translocation at SHL3 using an alternative crosslinking site at H2B residue 93, unlike ISW2. (G) SWR1C does not induce any translocation at SHL3.5 of the guide strand complement to the template strand that SWR1C is predicted to bind. (H) DNA-histone mapping detecting SWR1C dimer exchange on the linker-proximal side of an asymmetric nucleosome.

**Table S1 Single-molecule FRET events, SWRIC eviction of H2A from the nucleosome**

| [ATP] 100 uM | | | | | | | | | |
|---|---|---|---|---|---|---|---|---|---|
| Type of Events | Tethered Nucleosomes (#) | Priming Events (#) | Priming tau* (s) | 95 % C.I. (s) | Median eviction (s) | Stdev (s) | Release Events (#) | Median Release (s) | Stdev (s) |
| Distal First | 939 | 164 | 51 | 44.1-60.0 | 2.5 | 2.1 | NAN | NAN | NAN |
| Proximal second | 939 | 69 | 90 | 72.3-116.2 | 2.5 | 1.8 | 69 | 65 | 32.0 |
| Proximal First | 939 | 40 | 36 | 27.1-50.5 | 2 | 1.0 | NAN | NAN | NAN |
| Distal second | 939 | 10 | 71 | 41.4-147.45 | 3.5 | 0.9 | 10 | 18 | 5.8 |
| Single Events | 939 | 352 | 51 | 46.6-57.4 | 2.5 | 1.5 | 352 | 63 | 21.2 |
| Aggregate Events | 939 | 635 | 55 | 51.0-59.6 | 2 | 0.6 | 431 | 96[#] | 68.0-120.5[##] |
| **[ATP] 5 uM** | | | | | | | | | |
| Type of Events | Tethered Nucleosomes (#) | Priming Events (#) | Priming tau* (s) | 95 % C.I. (s) | Median eviction (s) | Stdev (s) | Release Events (#) | Median Release (s) | Stdev (s) |
| Distal First | 940 | 156 | 73 | 62.7-85.9 | 3 | 1.7 | NAN | NAN | NAN |
| Proximal second | 940 | 73 | 112 | 90.4-143.2 | 2.5 | 2.1 | 73 | 88 | 51.5 |
| Proximal First | 940 | 15 | 39 | 24.7-69.3 | 2 | 2.2 | NAN | NAN | NAN |
| Distal second | 940 | 10 | 99 | 54.7-228.4 | 3 | 3.5 | 8 | 25 | 16.7 |
| Single Events | 940 | 315 | 64 | 58.2-76.6 | 3 | 2.2 | 315 | 92 | 61.0 |
| Aggregate Events | 940 | 569 | 73 | 67.2-79.3 | 3 | 0.6 | 398 | 136[#] | 114.0-166.5[##] |
| **[ATP] 0.5 uM** | | | | | | | | | |
| Type of Events | Tethered Nucleosomes (#) | Priming Events (#) | Priming tau* (s) | 95 % C.I. (s) | Median eviction (s) | Stdev (s) | Release Events (#) | Median Release (s) | Stdev (s) |
| Distal First | 865 | 85 | 70 | 57.5-88.1 | 3.5 | 2.0 | NAN | NAN | NAN |
| Proximal second | 865 | 34 | 137 | 100.4-197.2 | 2.5 | 1.1 | 34 | 112 | 67.7 |
| Proximal First | 865 | 13 | 70 | 43.4-131.5 | 2.5 | 2.4 | NAN | NAN | NAN |
| Distal second | 865 | 4 | 139 | 63.5-511.1 | 3 | 1.3 | 4 | 88 | 47.2 |
| Single Events | 865 | 189 | 94 | 82.1-109.2 | 2.5 | 4.4 | 189 | 134 | 125.0 |
| Aggregate Events | 865 | 325 | 92 | 82.8-102.9 | 2 | 0.7 | 227 | 219[#] | 204.5-240[##] |
| **[ATP] 0 uM** | | | | | | | | | |
| Type of Events | Tethered Nucleosomes (#) | Priming Events (#) | Priming tau* (s) | 95 % C.I. (s) | Median eviction (s) | Stdev (s) | Release Events (#) | Median Release (s) | Stdev (s) |
| Distal First | 962 | 66 | 132 | 105.2-170.7 | 4 | 2.6 | NAN | NAN | NAN |
| Proximal second | 962 | 2 | 151 | 54.2-1246.9 | 3.5 | 0.0 | 2 | 74 | 73.5 |
| Proximal First | 962 | 22 | 54 | 37.1-86.5 | 3 | 3.8 | NAN | NAN | NAN |
| Distal second | 962 | 0 | NaN | NaN | NaN | NaN | 0 | NaN | NaN |
| Single Events | 962 | 15 | 90 | 57.9-162.3 | 2.5 | 2.0 | 15 | 138 | 150.0 |
| Aggregate Events | 962 | 147 | 110 | 94.9-131.3 | 3 | 2.7 | 17 | 179[#] | 149.5-331[##] |

*tau = lifetime exponential fit of distribution, C.I. = 95 % confidence interval of exponential fit, [#]Half-life estimate Kaplien-Mier survival curve, [##]Half-life 95 % confidence interval

**Table S2. Unwrapping events during priming phase**

| | # Nucleosomes | #events | Freq.(sec^-1) | mean dwell (sec) |
|---|---|---|---|---|
| [ATP] 0 uM | 491 | 152 | 0.0016 | 1.49 +/- 2.32 |
| [ATP] 100 uM | 487 | 234 | 0.0038 | 3.4 +/- 5.8 |

**Table S3. Single-molecule colocalization of H2AZ-Cy3b with SWR1C-nucleosome**

| # tethered nucleosomes | #events | time for 50% bound (s) | alpha* (C.I. 95%) | Dwell tau1* (C.I. 95%) (s) | Dwell tau2* (C.I. 95%) (s) | <Dwell>** (s) | # Depo. events | median arrival before Depo. (s) | Delay between binding and deposition (s) |
|---|---|---|---|---|---|---|---|---|---|
| 307 | 1136 | 10 | 0.78 (0.73-0.82) | 11 (9.7-11.9) | 117 (91.0-155.9) | 34 | 29 | 75+/-40 | 38+/-20 |

*Values correspond to output from fitting dwelltime distribution to a double exponential, alpha represents fraction of tau1, 95% Confidence interval obtained from bootstrap analysis. **corresponds to average dwelltime = alpha*tau1+(1-alpha)*tau2

**Table S4 Oligonucleotides**

| | |
|---|---|
| oAM200 | 5' /5Biosg/CCAGTTACCTTCGGAAAAAGAGTTcagtgcttggtagtcGATCTCAACAGCGGTAAGATCC |
| oTG415 | 5' GGATCT/iCy3N/ACCGCTGTTGAGATC |
| oTG416 | 5' AACTCT/iCy5N/TTTCCGAAGGTAACTGG |
